# Telomere-to-telomere gapless chromosomes of banana using nanopore sequencing

**DOI:** 10.1101/2021.04.16.440017

**Authors:** Caroline Belser, Franc-Christophe Baurens, Benjamin Noel, Guillaume Martin, Corinne Cruaud, Benjamin Istace, Nabila Yahiaoui, Karine Labadie, Eva Hřibová, Jaroslav Doležel, Arnaud Lemainque, Patrick Wincker, Angélique D’Hont, Jean-Marc Aury

## Abstract

Long-read technologies hold the promise to obtain more complete genome assemblies and to make them easier. Coupled with long-range technologies, they can reveal the architecture of complex regions, like centromeres or rDNA clusters. These technologies also make it possible to know the complete organization of chromosomes, which remained complicated before even when using genetic maps. However, generating a gapless and telomere-to-telomere assembly is still not trivial, and requires a combination of several technologies and the choice of suitable software. Here, we report a chromosome-scale assembly of a banana genome (*Musa acuminata*) generated using Oxford Nanopore long-reads. We generated a genome coverage of 177X from a single PromethION flowcell with near 17X with reads longer than 75Kb. From the 11 chromosomes, 5 were entirely reconstructed in a single contig from telomere to telomere, revealing for the first time the content of complex regions like centromeres or clusters of paralogous genes.

## Introduction

Long-read technologies are now the standard for generating high-quality assemblies, especially for complex genomes such as plant genomes^1–4^. Although the impact of these technologies is undeniable, they still lack the maturity to reconstruct complete chromosomes from telomere to telomere. Generally, assemblies based on long-reads are complemented with long-range data, like optical maps or chromosomal conformation sequencing. Recently, The Telomere-to-Telomere (T2T) consortium proposed a telomere-to-telomere assembly of the X chromosome of the human genome^5^. This high-quality assembly of the human genome was based on a combination of several existing technologies: Oxford Nanopore technology (ONT), Pacific Biosciences (PACBIO), 10X Genomics (10X) and Bionano Genomics (BNG). Even if the final assembly is very contiguous, there are still several gaps, and the complete X chromosome was obtained by manual curation. This huge effort is not possible for all genome projects because it is far too expensive and time consuming. It is clear that these multilayer assemblies reveal the architecture of complex regions as well as the complete organization of chromosomes, which remained complicated before. Long-range technologies make it possible to organize contigs based on long-reads but they are not able to fill the gaps between these contigs. Indeed, usually complex regions like centromeres or telomeres still contain many gaps, depending on their repetitive content.

We selected the banana genome, a medium-size genome in the plant lineage (∼500 Mb), and hypothesized that recent improvement of the ONT technology, coupled with dedicated DNA extraction protocol and efficient software enable the reconstruction of gapless and Telomere-to-Telomere chromosomes.

Banana species are monocotyledonous plants and part of the Zingiberales order and of the Musaceae family. Bananas are mostly cultivated in tropical and subtropical countries, and their fruits are the basis of the diet of several hundred million people and are massively exported to industrialized countries. Four genetic groups have been predicted to be involved in the origins of cultivars, mainly through inter(sub)specific hybridization and with different extents of contribution : *Musa acuminata* including various subspecies (A-genome), *Musa balbisiana* (B-genome), *Musa Schizocarpa* (S-genome) and species of the *Australimusa* section (T-genome). Two events appeared during banana domestication: the transition from wild to edible diploids and the emergence of triploids from edible diploids^6–8^. Recent results suggest that edible cultivar origins are more complex than expected, involving multiple hybridization steps, resulting in inter(sub)specific mosaic genomes. They also revealed that additional genetic pools to the ones expected were involved, for which the wild contributors are still unidentified^6^. In addition, large structural variations in form of reciprocal translocations and a few inversions have been characterized in genetic pools involved in cultivar origins and found widespread in cultivated germplasm^6^. The complexity of these genomes underlies the importance of producing high-quality assemblies of banana genomes to decipher their evolutionary history and to support genetic studies.

In this context, two versions of the *Musa acuminata* ‘DH-Pahang’ genome have already been proposed^9,10^. The first draft version of the genome (V1) was published in 2012 and based on 454, Sanger (fosmids and BAC-ends) and Illumina sequencing. Furthermore, scaffolds were organized using a sparse genetic map, resulting in the anchoring of 63% of the estimated genome size. The second version (V2) was published in 2016 and added illumina long-insert sequences, a low-contiguity optical map as well as a more dense genetic map. Martin et al. proposed an assembly of the 11 chromosomes that included 76% of the estimated genome size. Herein we propose to generate a new version (V4) of the DH-Pahang assembly based on nanopore long-reads.

## Results

### Highly continuous genome assembly of the banana genome

The efficiency of long-reads sequencing depends on the quality of the DNA extraction. Here, DNA was extracted following a plant-dedicated protocol provided by Oxford Nanopore Technologies (“High molecular weight gDNA extraction from plant leaves’’). This protocol was particularly effective and allowed us to obtain long DNA fragments (> 50 Kb). Residual short fragments were filtered out using the Short Read Eliminator (SRE) XL kit (Circulomics, MD, USA). Almost 93Gb of nanopore sequences were obtained with a single PromethION flowcell R9.4.1. The 5.2M reads had a N50 of 31.6Kb and the genome was covered at 17X with reads longer than 75Kb. This high-quality set of long reads was assembled using several bioinformatics tools and the assembly obtained with NECAT^11^ was retained. This assembly was polished first using long reads with Racon^12^ and Medaka^13^ and then using Illumina short-reads with Hapo-G^14^. This assembly, based on nanopore long reads and without long-range information, was composed of 124 contigs (larger than 50Kb) and had a cumulative size of 485Mb. Half of the assembly size was composed of contigs larger than 32Mb and only 17 contigs covered 90% of the total length (Table 1). More importantly the seven largest contigs had a size compatible with complete chromosomes (ranging from 47.7 to 32.1 Mb). The anchoring of contigs was performed following methodology described in Martin et al^10^. As expected, the five largest contigs correspond to complete chromosomes and harbor telomeric repeats at both extremities (Figure 1). The six remaining chromosomes were composed of a small number of contigs (between four and eight). Unsurprisingly, the remaining gaps are mainly located in rDNA clusters: 5S for chromosomes 1,3 and 8 and 45S for chromosome 10 or in other tandem and inverted repeats: chromosome 1 and 5 (Figure 2A). These rDNA clusters are composed of a large number of tandemly repeated genes and are generally very difficult to assemble. Even if these clusters still contain few gaps, it is now possible to decipher the architecture of these large and complex regions. In addition, smaller contigs not anchored to the 11 chromosomes correspond to the chloroplastic and mitochondrial genomes (one and 45 contigs, respectively). A total of 37 contigs were filtered out because they were included in larger contigs and contained highly repeated sequences.

**Table 1:** Comparison of *Musa acuminata* (DH-Pahang) genome assemblies.

|  | D'hont et al.<sup>9</sup><br>V1 | Martin et al.<sup>10</sup><br>V2 | This study<br>V4 |
| --- | --- | --- | --- |
| Number of contigs | 29,437 | 19,312 | 124 |
| Cumulative size (bp) | 390,477,446 | 405,522,014 | 484,058,756 |
| N50 (bp) | 28,319 | 43,237 | 32,091,396 |
| L50 | 3,428 | 2,363 | 7 |
| N90 (bp) | 5,108 | 9,026 | 6,704,534 |
| L90 | 16,551 | 10,327 | 16 |
| Longest contig (bp) | 306,660 | 602,020 | 47,719,527 |
| Number of chromosomes | 11 | 11 | 11 |
| Cumulative size (bp) | 331,812,599 | 397,008,016 | 468,821,802 |
| Cumulative size (ACGT only) | 286,824,765 | 363,519,833 | 468,133,046 |
| N50 (bp) | 30,470,408 | 37,593,364 | 43,931,232 |
| L50 | 5 | 5 | 5 |
| N90 (bp) | 25,514,024 | 29,070,452 | 34,826,100 |
| L90 | 10 | 10 | 10 |
| Longest (bp) | 35,439,739<br>A08 | 44,889,171<br>A08 | 51,314,288<br>A08 |
| % of N | 13.56% | 8.44% | 0.68% |
| % of estimated genome | 63.4% | 75.9% | 89.6% |
| % of estimated genome (ACGT only) | 54.8% | 69.5% | 89.5% |
| Complete BUSCO (N=1,614) | 92.6% | 98.5% | 98.8% |

**Figure 1:**
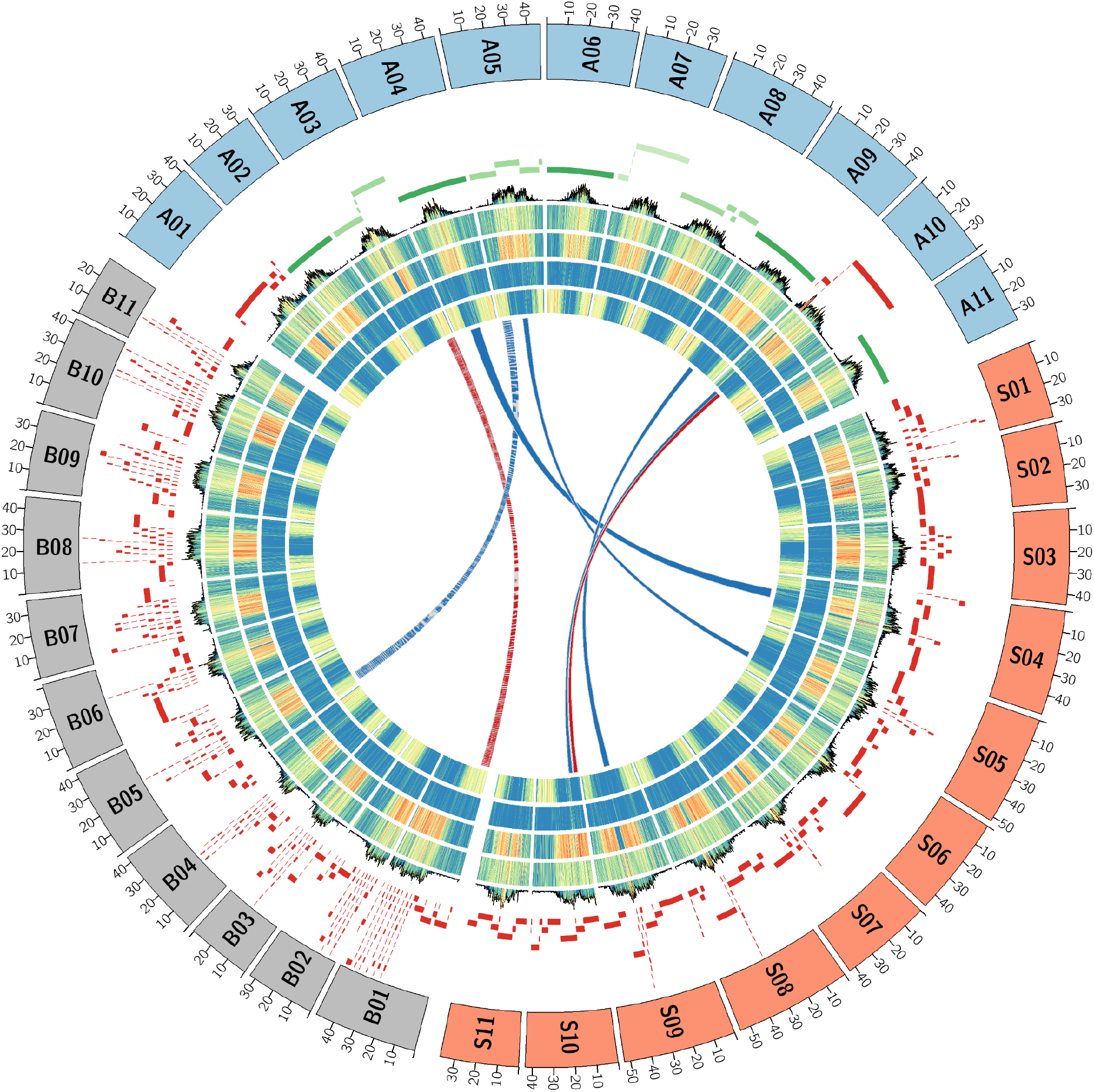
Musa genomes architecture comparison. The tracks represent the following elements (from outer to inner) : (1) Schematic representation of *M. acuminata* (A), *M. balbisiana* (B) and *M. schizocarpa* (S) chromosomes, (2) contigs colored in green if the chromosome is composed of 1 to 4 contigs, in red if the chromosome is composed of more than 5 contigs. (3) density of the centromeric repeats. (4) density of the Gypsy elements. (5) density of the Copia elements. (6) density of the DNA transposons. (7) density of genes. (8) synteny relationships. The red lines show translocations between B01 and A03 and between S10 and A10. The blue lines show inversions between B05 and A05, S04 and A04, S05 and A05, S09 and A09.

**Figure 2:**
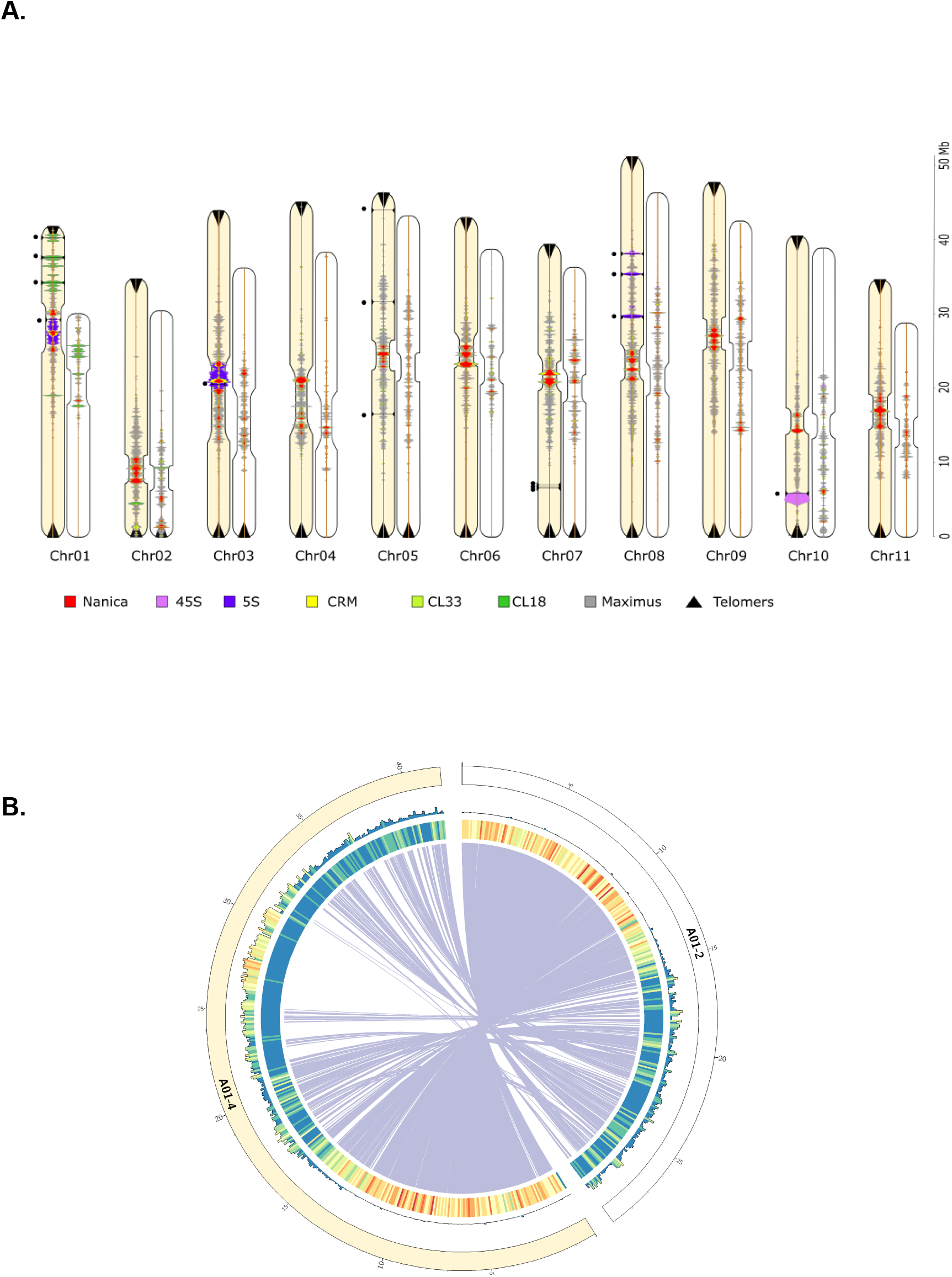
Comparison of the V2 and V4 assemblies. **A**. Localization and density of several repeated elements on chromosomes of the V2 (light orange) and V4 (white) assemblies (scale in Mb on the right) with Nanica LINE (Red), CRM chromovirus Gypsy retrotransposon (yellow), 5S rDNA (Blue), 45S rDNA (violin), tandem repeat cluster CL18 (dark green), tandem repeat cluster CL33 (light green), Maximus Copia retrotransposon (Grey) and telomeric sequences (black triangles). Horizontal black lines and black dots correspond to the 15 remaining gaps in the V4 assembly. **B**. Comparison of the A01 chromosomes of the V2 and V4 assemblies. Tracks represent the following elements (from outer to inner) : (1) density of the centromeric repeats. (2) density of genes. (3) synteny relationships between the V2 chromosome 1 and the V4 chromosome 1.

### Validation of telomere-to-telomere chromosomes

A kmer analysis and a first alignment of the largest contigs with the previous version of the DH-Pahang assembly did not reveal chimeric contigs (Figures S4 and S5). In addition, all eleven chromosomes harbor T3AG3 repeat at both sides, underlining the complete assembly of chromosome ends.

However, we decided to validate the quality of our assembly using two Bionano optical maps that were generated using the Saphyr instrument commercialized by Bionano Genomics (BNG). High molecular weight DNA was extracted and labeled with two different enzymes (DLE-1 and BspQI). The DLE-1 and BspQI optical maps were 469Mb and 474Mb length respectively and had a N50 of 35Mb and 16Mb respectively. We used these two optical maps to first validate the contigs, and then order and orient them. As a result, only one contig of 380Kb, composed of tandem repeated elements, was flagged as conflictual with the optical maps and split into two contigs (Figure S6). All other contigs were in accordance with the maps, which strongly validate the accuracy of the NECAT assembler. The 124 contigs were ordered in 96 scaffolds using the Bionano Solve workflow and the BiscoT^15^ software (88 scaffolds correspond each to one contig). In the end, eight of the eleven chromosomes are represented by a single scaffold and the other four remain in two scaffolds. The whole genome assembly contains only fifteen gaps that are concentrated in large highly repetitive regions (Table S6).

### Comparison of *Musa acuminata* assemblies

Unsurprisingly, compared to previous versions, the contiguity of our DH-Pahang assembly is greatly improved. The contig N50 goes from a few tens of Kb (28 Kb and 43 Kb for V1 and V2 respectively) to a few tens of Mb (32 Mb). More importantly, the cumulative size is closer to the estimated genome size, suggesting that complex regions are better represented in this new release (Table 1). With a very small number of contigs, anchoring on the eleven chromosomes using the genetic map was easier especially in the centromeric regions which are generally difficult to organize due to their lower density of genetic markers. The size of the DH-Pahang genome was estimated by flow cytometry at 523 Mb^9^. The 11 chromosomes of our long read assembly cover almost 90% of this estimated size, while the first two versions were largely incomplete (55% and 70% respectively, Table 1). As a consequence, the assembled size of each chromosome has increased, between 7% for chromosome 10 to 43% for chromosome 1 (Figure 2A and Figure S5). Chromosomes of the two versions were aligned and large genomic regions (>100Kb) absent in the previous assembly were reported. The 11 chromosomes totalized 247 new regions that covered 141.4 Mb, i.e. 29.2% of the assembly (Figure S9). The largest region, close to 6 Mb, is localized on chromosome 1. Unsurprisingly, these blocks are mainly composed of repeated elements (more than 85%), and localized in centromeric regions or rDNA clusters (Figure S10).

We annotated 246 Mb of the genome (52.6%) as transposable elements (TE), compared to 152 Mb in V2, which illustrates the much better representation and completion of these repetitive elements in the V4 assembly (Table S8 and Figure S8). Figure 2A shows the distribution of several tandem repeats and TE along the chromosomes, including the Maximus Copia retrotransposons which are the most abundant TEs in the Pahang genome.

### Architecture of centromeric regions and rDNA clusters

Earlier cytogenetic analysis showed that a long interspersed element (LINE), named Nanica, is present in the centromeric regions of banana^9,16^. Very few LINE sequences were present in the first release of the assembly despite being present in unassembled reads^9^ and they had a scattered distribution on the pericentromeric and centromeric regions of the V2 assembly. In this new assembly, clusters of Nanica tandem repetitions are found grouped in the centromeric regions of all chromosomes (Figures 2A and S8). Several elements of chromovirus CRM clade, a lineage of Ty3/Gypsy retrotransposons, were also found restricted to these centromeric regions. Some members of this plant retroelement have been shown to have the ability to target their insertion almost exclusively to the functional centromeres^17,18^. The position of two other tandem repeats (CL18 and CL33) previously identified^16^ could also be refined between V2 and V4 and the localization of the main clusters on chromosome 1 and 2 are in accordance with cytogenetic karyotypes^16^.

Regarding the 5S rDNA sequences, in the V2 assembly, they were present in a few numbers in chromosomes 5, 9 and 8 spanning 7,5 kb of sequences (around 130 gene units) (Table S10). In the new assembly, six major loci containing 5S rDNA gene clusters are present accounting for around 7,696 gene units. Three clusters are located on one arm of chromosome 8 representing in total around 2.2 Mb and two large clusters of 5S rDNA repeat have been integrated to chromosome 1 and 3 centromeric regions, representing around 3.5Mb and 4Mb, respectively. These results are in accordance with previous cytogenetic results^16^ and the position of the rDNA clusters have been clarified showing that they colocalize with the centromeric Nanica clusters of these chromosomes. Clusters of 5S rDNA are organized in canonical gene/spacer tandem repeat of different length due to the insertion in the spacer of various repeated elements such as Nanica or CRM sequences as observed in the 5S cluster of the centromeric regions of chromosomes 1 and 3 (Figure 3).

**Figure 3:**
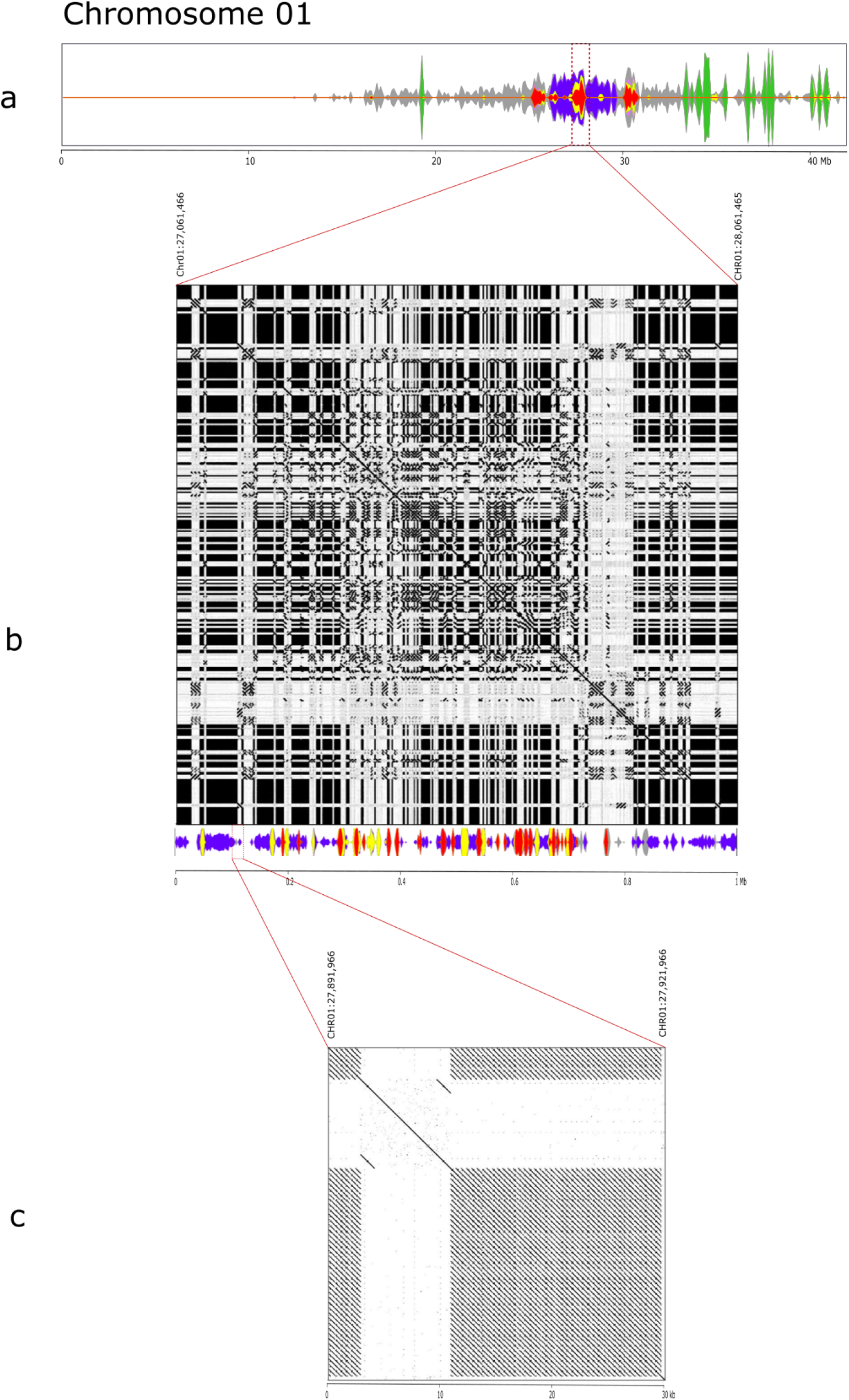
Fine structure and density of main (peri)centromeric repeated sequences on chromosome. **1** with Nanica LINE (Red), CRM chromovirus Gypsy retrotransposon (yellow), 5SrDNA (Blue), tandem repeat cluster CL18 (dark green), Maximus Copia retrotransposon (Grey) : **a**. entire chromosome 1: **b**. zoom and dot-plot alignment of a 1Mb segment in the centromeric region containing Nanica, 5SrDNA and CMR repeats, **c**. zoom and dot-plot alignment of a 30Kb segment containing 5S rDNA repeats and a CMR.

Furthermore, a large cluster of 1.8 Mb containing around 110 45S rDNA units, consisting in canonical gene/spacer tandem repeat, is localized on chromosome 10 between positions 4.4 and 6.2Mb (Figure 2A).

### Tandemly duplicated genes

Gene duplication is an important evolutionary mechanism that contributes to the appearance of novel functions and to adaptation. The events leading to gene duplication have contributed to important plant agronomic traits, such as grain quality, fruit shape, and flowering time^19^. A special case of gene duplication relates to genomic/tandem duplication events which generate, locally, repetitive regions in the genome. These tandemly duplicated genes (TDGs) are generally harder to capture in short-read assemblies, especially in the case of recent multi-copy clusters. We found 1,700 genes that have been annotated only in our long-read assembly (Table S9 and Figure S7). These genes are distributed over the different chromosomes, with chromosomes 1 and 10 having the greatest number of new genes, 13.3% and 14.3% respectively (Table S9). Interestingly, a large proportion (38.3%) of these genes are TDGs included in a gene cluster and the proportion of TDGs in new genes is higher when compared to the whole gene catalogue (38.3% versus 9.9%).

By focusing on TDGs and detecting gene clusters in the short and long-read assemblies, we found 31% more clusters in the V4 compared to V2 assembly (1,134 compared to 866 clusters). These blocks of TDGs contain respectively 3,649 and 1,134 genes. The largest in the long-read assembly contains 38 genes on chromosome 7 (between 31.8 and 32.8Mb) and was split into two smaller clusters of 11 and 9 genes in the V2. This TDG cluster is located in a region with several gaps in previous versions, and we found three regions (165Kb, 111Kb and 110Kb) between positions 31.8 and 32.3Mb of the chromosome 7 that are specific to the V4 assembly. These new regions allow the creation of a complete cluster of TDGs (Figure 4A) which contain motifs of the terpene synthase family that are responsible for the synthesis of terpenoid compounds playing a role in plant flavor^20^ and more generally in the interactions between the plant and its environment^21,22^. This family is known to contain TDGs and is expanded in several plant species^23^.

**Figure 4:**
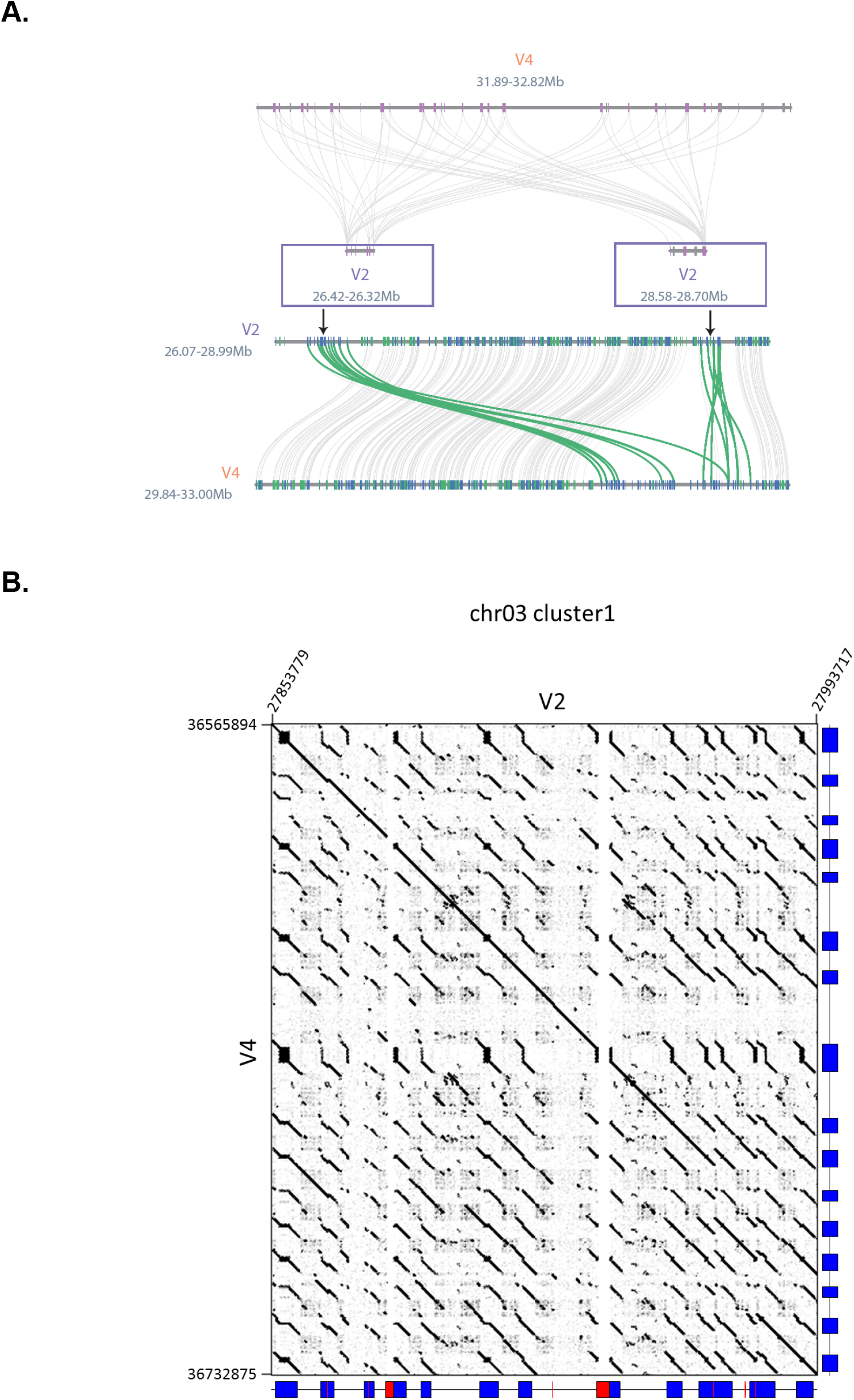
Comparison of tandemly duplicated gene regions in the V2 and V4 assemblies. **A**. Gene clusters of the terpene synthase family. The cluster is located between 31.89 and 32.82Mb on the V4 chromosome 7. The top figure represents the correspondence of terpene synthase genes between the two versions. The bottom figure represents the synteny of all genes in the same region. Terpene synthase genes synteny relationships are colored in green. Genes are colored according to their orientation (blue if forward and green otherwise). **B**. Comparison of the structure of a NLR cluster on chromosome 3. The predicted NLR loci for each version are represented on the right side and at the bottom of the dot plots by blue boxes. Red boxes represent regions bearing undetermined nucleotides. Region coordinates are also indicated.

### Resistance Genes

Plant disease resistance genes encoding proteins with nucleotide-binding leucine-rich repeat (NLR) domains are often clustered in genomes, sometimes forming large, rapidly evolving clusters of highly homologous genes^24^. The NLR-annotator program^25^ allows the identification of NLR loci i.e. genomic regions likely associated with an NLR gene (or pseudogene). A total of 128 NLR loci were detected in this assembly compared to 111 loci in the V2 assembly (Table 2, S12 and S13). Four major clusters of NLR loci were found in this assembly: two in chromosome 3, one in chromosome 7, one in chromosome 10 (Figures 4B and S13). They all have a larger size compared to V2, with sizes ranging from 132Kb up to 227Kb and additional detected NLR loci. These clusters were improved in sequence quality with a complete absence of undetermined nucleotides within the corresponding genomic regions (Table S11).

**Table 2:** Comparison of gene prediction statistics.

| Reference | Martin et al.<sup>10</sup><br>V2 | This study<br>V4 |
| --- | --- | --- |
| # number of genes | 35,276 | 36,979 |
| #exons<br>per spliced gene<br>(avg:med) | 6.08 : 5 | 6.05 : 4 |
| Gene sizes<br>(avg:med) | 4,542 : 2,824 | 4,604.68 : 2,758 |
| CDS sizes<br>(avg:med) | 1,171 : 981 | 1,180 : 972 |
| Complete BUSCO<br>(N=1,614) | 98,5% | 98.8% |
| NLR loci | 111 | 128 |

### Comparison of A, B and S-genomes

The B and S genomes have already been recently sequenced using a long-read strategy^26,27^. Taking into account this new version of the *M. acuminata* genome, three high-quality banana genomes are now available. The A and S genomes were sequenced using ONT while the B genome was sequenced using the PACBIO technology. Interestingly, the two ONT assemblies have a higher contiguity (contig N50 of 32 Mb and 6.5 Mb compared to 1.8 Mb) suggesting the usage of longer reads, or the difficulty to extract and sequence long DNA fragments with the PACBIO device (Table 3). Indeed, the PACBIO library was size-selected in order to obtain fragments around 20Kb^27^, which is perhaps an optimal condition for PACBIO sequencing. The B genome was assembled from reads with an N50 of 16.6 Kb whereas A and S enomes were assembled with reads having a N50 of 31,6 Kb and 24.4 Kb respectively. As a consequence, chromosomes of the A and S genomes contain less gaps. The eleven A and S chromosomes contain 15 and 166 gaps respectively whereas B chromosomes contain 683 gaps and no chromosome is gapless (Figures 1). Centromeric regions, detected with centromeric repeats, are very fractionated in the case of the PACBIO-based assembly (from 24 contigs for the chromosome 7 to 111 contigs for the chromosome 1), underlying the importance of ultra long reads to resolve these highly repetitive regions.

**Table 3:**
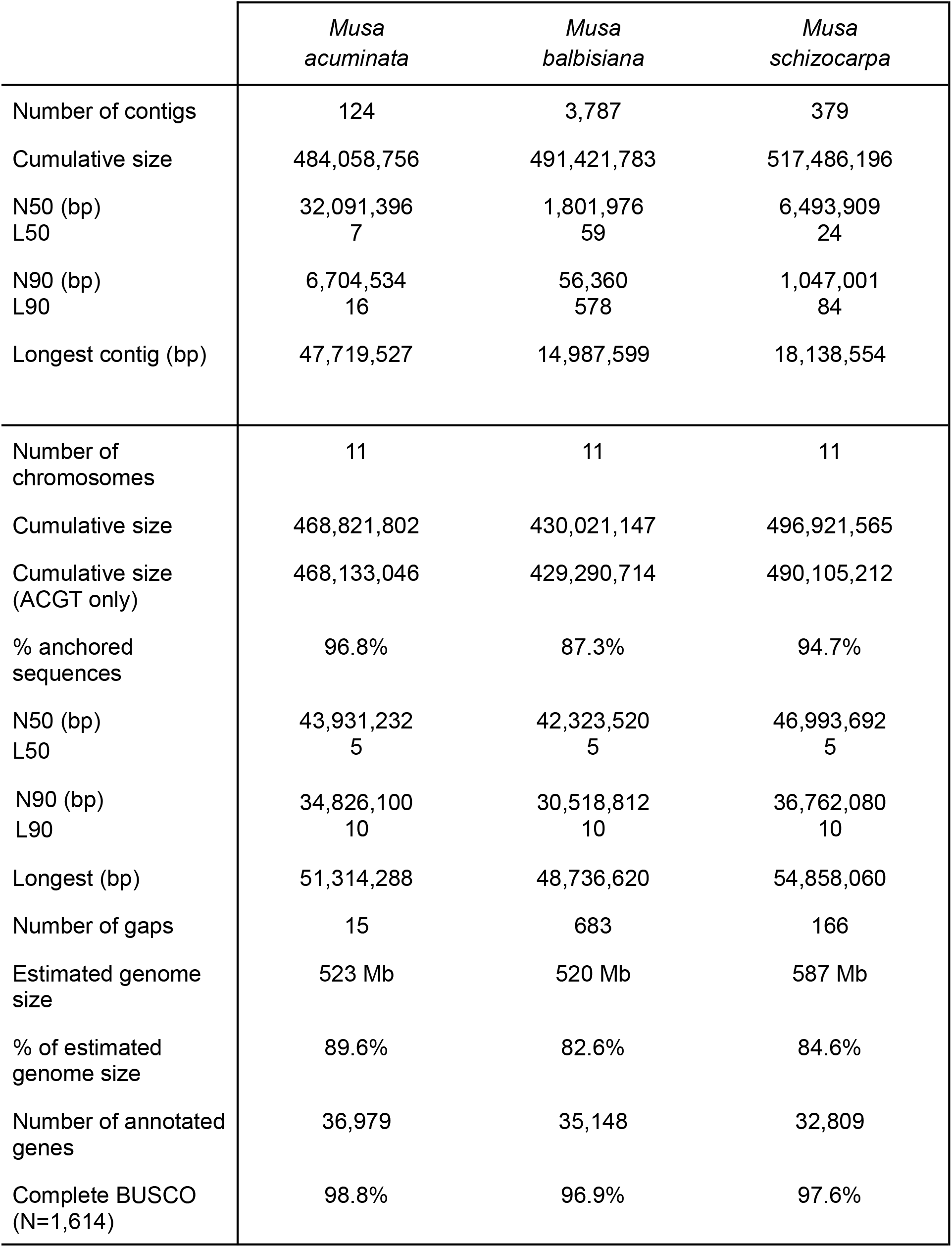
Comparison of A (*Musa acuminata*), B (*Musa balbisiana*) and S (*Musa schizocarpa*) genomes assemblies.

|  | *Musa*<br>*acuminata* | *Musa*<br>*balbisiana* | *Musa*<br>*schizocarpa* |
| --- | --- | --- | --- |
| Number of contigs | 124 | 3,787 | 379 |
| Cumulative size | 484,058,756 | 491,421,783 | 517,486,196 |
| N50 (bp) | 32,091,396 | 1,801,976 | 6,493,909 |
| L50 | 7 | 59 | 24 |
| N90 (bp) | 6,704,534 | 56,360 | 1,047,001 |
| L90 | 16 | 578 | 84 |
| Longest contig (bp) | 47,719,527 | 14,987,599 | 18,138,554 |
| Number of chromosomes | 11 | 11 | 11 |
| Cumulative size | 468,821,802 | 430,021,147 | 496,921,565 |
| Cumulative size (ACGT only) | 468,133,046 | 429,290,714 | 490,105,212 |
| % anchored sequences | 96.8% | 87.3% | 94.7% |
| N50 (bp) | 43,931,232 | 42,323,520 | 46,993,692 |
| L50 | 5 | 5 | 5 |
| N90 (bp) | 34,826,100 | 30,518,812 | 36,762,080 |
| L90 | 10 | 10 | 10 |
| Longest (bp) | 51,314,288 | 48,736,620 | 54,858,060 |
| Number of gaps | 15 | 683 | 166 |
| Estimated genome size | 523 Mb | 520 Mb | 587 Mb |
| % of estimated genome size | 89.6% | 82.6% | 84.6% |
| Number of annotated genes | 36,979 | 35,148 | 32,809 |
| Complete BUSCO (N=1,614) | 98.8% | 96.9% | 97.6% |

Overall, the synteny conservation between the three genomes is high, we detected one inversion between the chromosome B05 of *Musa balbisiana* and the chromosome A05 of *Musa acuminata* and a translocation between the chromosome B01 and the chromosome A03 as already reported^27^ (Figure 1 and Figure S11). Four inversions between *Musa schizocarpa* and *Musa acuminata* were also detected on the chromosomes S10, S04, S05 and S09 and one translocation on chromosome A10 (Figure 1 and Figure S12). The corresponding regions on chromosome A10 contain the 45S rDNAS gene cluster. The contigs organization in these five regions was manually validated in the two genome assemblies using optical maps.

## Discussion

Long read sequencing technology emergence have paved the way for high quality genome assemblies. The rapid evolution of the DNA extraction protocols now allows the community to sequence very long DNA sequences. Coupled with the evolution of the bioinformatic tools, the generation of high-quality assemblies has been greatly simplified. The latest improvements of the ONT technology, especially the base-calling efficiency, result in a decrease of the error rate. Several assembly tools were specially developed around long reads and are able to manage noisy reads and have their specificity as well as a specific margin of progress. We think that it is still important to use the latest release of several assemblers and choose the most efficient for each genome assembly project.

In this study, we combined recent development from DNA extraction, sequencing and genome assembly and showed that plant chromosomes can now be assembled in a single contig, gapless and from telomere to telomere, at least to a certain extent. We chose the genome of *Musa acuminata*, the first monocotyledonous species sequenced outside Poales, because its reference genome, even of low-quality, has been widely used and constitutes an important resource for the scientific community. To date, three *Musa* species: *Musa acuminata, Musa balbisiana* and *Musa schizocarpa* are available at the chromosome-scale. These three species are of particular interest because they are involved in the origin of banana cultivars, their high-quality genome assemblies will thus be a valuable resource to explore the evolutionary history and biology of current banana cultivar.

We generated a highly contiguous assembly of the eleven chromosomes of *Musa acuminata* of which five were obtained in a single contig. At the same time, optical maps were used to validate the nanopore assembly. It is important to mention that only one small contig, essentially composed of repetitive elements, was detected as potential chimera underlining the high quality of the contigs produced by the NECAT assembler. All eleven chromosomes, build with the help of a genetic map, contain telomeric repeats at both ends which is an important element in asserting on the one hand that the reconstruction of the chromosomes is of good quality and on the other hand that the still missing part of the genome is contained in the remaining fifteen gaps, although 7 are of unknown length. One of the advantages of optical maps is that the size of the gaps can be estimated if the map is sufficiently contiguous (Table S6 and Figure S14).

Comparison of the distribution of repeated sequences (tandem repeat and TE) between V2 and V4 showed that the integration of these elements that were typically difficult to assemble with past technologies are greatly improved in the new assembly and are now very congruent with cytogenetic karyotype. All centromeres are now clearly identified with large clusters of Nanica LINE tandem repeats and CMR TE, and in addition, for two of them large clusters of 5S rDNA tandem repeats. Such case of recruitment of 5S rDNA gene array in centromere was also reported in one of the switchgrass chromosomes^28^. This high-resolution of centromeric regions opens new avenues to study how satellites repeats originate and evolve in the centromeric region and more generally to better understand the organization and functioning of centromeres that are essential chromosomal domains for kinetochore assembly and correct chromosome segregation^29,30^. In addition, chromosome reciprocal translocations were recently shown to have accompanied subspecies evolution in *Musa*^6^, and some of them have their break points in centromeric regions. Having access to the sequence of these centromeric regions will permit investigating the mechanism and sequences involved in the origin of these translocations. Finally, comparison with the *Musa acuminata* V2 assembly highlights a higher proportion of each class of transposable elements, and a large amount of additional sequences in the centromeric regions, like Nanica elements, or large retro-transposon derivatives.

It is often mistakenly thought that short-read assemblies are complete at the gene level. This hypothesis is mainly based on the results given by the BUSCO software which only focus on single-copy genes. Accordingly, the gene content completeness was already high, in previous *M. acuminata* assembly versions, according to the BUSCO score. However, here, using ultralong reads, we were able to assemble many additional copies of tandemly duplicated gene (TDGs) clusters which contain important gene families like terpene synthases or disease resistance genes. Banana crops are currently particularly threatened by diseases including Black leaf streak disease that requires massive use of pesticide^31^ and by a new strain of Fusarium wilt (Tropical Race 4) that is currently spreading around the world and for which no chemical control is possible^32^. This new assembly will facilitate the search for resistance genes to these devastating diseases^33^.

Finally, we showed that gapless and telomere-to-telomere assembly of chromosomes is now possible thanks to long-read sequencing. The critical point remains the DNA extraction protocols that generally need adaptation for each species. These closed assemblies will allow new discoveries and will shed new light on these genomes in particular in complex repetitive regions such as centromeres which have essential biological function but are so far poorly characterized.

## Methods

### Plant material

Double haploid *Musa acuminata* spp *malaccensis* (*DH-Pahang*) plant material was obtained from the CRB Plantes Tropicales Antilles CIRAD-INRA Guadeloupe under the collection number PT-BA-00461.

### DNA extraction

For Illumina sequencing libraries, DNA was extracted using a modified mixed alkyl trimethyl ammonium bromide (MATAB) procedure^34^. A total of 2 g freshly harvested leaves was ground in liquid nitrogen with a mortar and pestle and immediately transferred to 12 ml of 74°C prewarmed extraction buffer containing 100 mM Tris-HCl, pH 8, 20 mM EDTA, 1.4 M NaCl, 2% w/v MATAB, 1% w/v PEG6000 (polyethylene glycol), 0.5% w/v sodium sulfite and 20 mgl^− 1^ RNAse A. Crude extracts were maintained for 20 min at 74°C, extracted with an equal volume of chloroform-isoamyl alcohol (24:1) and transferred to clean tubes. DNA was recovered by centrifugation after adding 10 ml isopropanol. DNA precipitates were briefly dried, washed with 2 ml of 70% ethanol and resuspended in 1 ml sterile water. Extract quality was evaluated using pulse field gel electrophoresis for size estimation and spectrophotometry (A260/A280 and A260/A230 ratios) for purity estimation. DNA samples with a fragment size above 50 kb, a A260/A280 ratio close to 2 and a A260/A230 ratio above 1.5 were kept.

In order to generate long reads on the Oxford Nanopore Technologies devices, high-quality and high-molecular-weight DNA is needed. To that end, DNA was isolated following the protocol provided by Oxford Nanopore Technologies, “High molecular weight gDNA extraction from plant leaves” downloaded from the ONT Community in March, 2019 (CTAB-Genomic-tip). This protocol involves a conventional CTAB extraction followed by purification using the commercial Qiagen Genomic tip (QIAGEN, MD, USA), but size selection was performed using Short Read Eliminator XL (Circulomics, MD, USA) instead of AMPure XP beads. Briefly, 1,2g of leaves were cryoground in liquid nitrogen. The fine powder was transferred to 20 mL of Carlson buffer (100 mM Tris-HCl pH 9.5, 2% CTAB, 1.4 M NaCl, 1% PEG 8000, 20 mM EDTA, 0.25% b-mercaptoethanol (v/v)) prewarmed to 65 °C. Then 40 µl of RNase A (100 mg/ml) was added before incubation at 65°C for 1h (with intermittent agitation). Proteins removal was performed by addition of one volume of chloroform and centrifugation at 5,500 g for 10 minutes at 4 °C. DNA was then precipitated with 0,7V of isopropanol and centrifugation at 5,500 g for 30 minutes at 4 °C. The pellet was then purified using the Qiagen Genomic tip 100/G, following the manufacturer’s instruction: DNA pellet was first dissolved at 50°C for 15 min in 9,5 mL of G2 buffer before loading onto the pre-equilibrated Genomic tip column. Purified gDNA was finally precipitated with 0.7 volumes of isopropanol, washed with 2 ml of 70% ethanol, dried, and eluted in 100 µL of TE Buffer. DNA was quantified by a dsDNA-specific fluorometric quantitation method using Qubit dsDNA HS Assays (ThermoFisher Scientific, Waltham, MA). DNA quality was checked on a 2200 TapeStation automated electrophoresis system (Agilent, CA, USA) (Figure S1).

Generating optical maps requires high molecular weight (HMW) DNA. Here HMW DNA of *M. acuminata DH Pahang* was prepared according to Safář et al^35^ with several modifications. Briefly, 0.5 cm long segments of leaf midribs and young leaf tissues were fixed for 20 min at 4°C in Tris buffer (10 mM Tris, 10 mM EDTA, 100 mM NaCl, pH 7.5) containing 2% formaldehyde. After three 5-min washes in Tris buffer, the segments were homogenized using chopping by a razor blade in petri dish containing 1 ml of ice-cold IB buffer (15mM Tris, 10mM EDTA, 130mM KCl, 20mM NaCl, 1mM spermine, 1mM spermidine and 0.1 % Triton X-100, pH 9.4) and immediately before use, 33 μl of β-mercaptoethanol were added to 10 ml of IB buffer. Nuclei suspension was passed through a 50-μm nylon mesh and stained with DAPI at a final concentration of 2 μg/mL. Six batches of 900,000 G1-phase nuclei were sorted into 77 µl of IB buffer with β-mercaptoethanol in 1.5 ml polystyrene tubes using a FACSAria SORP flow cytometer and sorter (Becton Dickinson, San José, CA, United States) equipped with solid-state UV laser. One 20 µl agarose mini plug was prepared from each batch of nuclei according to Šimková et al^36^. Miniplugs were washed and solubilized using Agarase Enzyme (Thermo Fisher Scientific) to release high molecular weight (HMW) DNA. HMW DNA was further purified by drop dialysis and was then homogenized a few days prior to the quality control.

The concentration and purity of the extracted DNA was evaluated using a Qubit fluorometer (Thermo Fisher Scientific) and a Nanodrop spectrophotometer (Thermo Fisher Scientific). DNA integrity was checked by pulsed field gel electrophoresis (Pippin Pulse, Sage Science). DNA molecules were detectable between 50Kb to 300Kb in size.

### Illumina PCR-Free library preparation and sequencing

DNA (1.5 μg) was sonicated to a 100–1500-bp size range using a Covaris E220 sonicator (Covaris, Woburn, MA, USA). The fragments were end-repaired and 3′-adenylated. Illumina adapters were added using the Kapa Hyper Prep Kit (KapaBiosystems, Wilmington, MA, USA). The ligation products were purified with AMPure XP beads (Beckmann Coulter Genomics, Danvers, MA, USA). The libraries were quantified by qPCR using the KAPA Library Quantification Kit for Illumina Libraries (KapaBiosystems), and the library profiles were assessed using a DNA High Sensitivity LabChip kit on an Agilent Bioanalyzer (Agilent Technologies, Santa Clara, CA, USA). The libraries were sequenced on an Illumina HiSeq2500 instrument (Illumina, San Diego, CA, USA) using 250 base-length read chemistry in paired-end mode.

After the Illumina sequencing, an in-house quality control process was applied to the reads that passed the Illumina quality filters. The first step discards low-quality nucleotides (Q < 20) from both ends of the reads. Next, Illumina sequencing adapters and primer sequences were removed from the reads. Then, reads shorter than 30 nucleotides after trimming were discarded. These trimming and removal steps were achieved using in-house-designed software based on the FastX package^37^. The last step identifies and discards read pairs that are mapped to the phage phiX genome, using SOAP aligner^38^ and the Enterobacteria phage PhiX174 reference sequence (GenBank: NC_001422.1). This processing, described in Alberti et al^39^, resulted in high-quality data.

### MinION and PromethION library preparation and sequencing

The library was prepared according to the following protocol, using the Oxford Nanopore SQK-LSK109 kit. Genomic DNA fragments (4µg) were repaired and 3’-adenylated with the NEBNext FFPE DNA Repair Mix and the NEBNext® Ultra^™^ II End Repair/dA-Tailing Module (New England Biolabs, Ipswich, MA, USA). Sequencing adapters provided by Oxford Nanopore Technologies (Oxford Nanopore Technologies Ltd, Oxford, UK) were then ligated using the NEBNext Quick Ligation Module (NEB). After purification with AMPure XP beads (Beckmann Coulter, Brea, CA, USA), half of the library was mixed with the Sequencing Buffer (ONT) and the Loading Bead (ONT) and loaded on a PromethION R9.4.1 flow cell. The second half of the library was loaded on the flow cell after a Nuclease Flush using the Flow Cell Wash Kit EXP-WSH003 (ONT) according to the Oxford Nanopore protocol. Reads were basecalled using Guppy version 4.0.1. The nanopore long reads were not cleaned and raw reads were used for genome assembly.

### Optical mapping

The Direct label and stain (DLS) labelling (DLE-1) and the Nick Label Repair and Stain (NLRS) labelling (BspQI) protocols were performed according to Bionano Genomics with 750ng and 600ng of DNA respectively. The Chip loadings were performed as recommended by Bionano Genomics.

### Long reads-based genome assembly

We generated three samples of reads : all reads, 30X of the longest reads and 30X of the filtlong^40^ highest-score reads. We then applied four different assemblers, Smartdenovo^41^, Redbean^42^, Flye^43^ and NECAT^11^ on these three subsets of reads (Table S1), with the exception of NECAT being only launched with all reads, as it applies a downsampling algorithm in its pipeline. Smartdenovo was launched with -k 17, as advised by the developers in case of larger genomes and -c 1 to generate a consensus sequence. Redbean was launched with ‘-xont -X5000 -g450m’ and Flye with ‘-g 450m’. NECAT was launched with a genome size of 450Mb and other parameters were left as default. After the assembly phase, we selected the best assembly (NECAT with all reads) based on the cumulative size and contiguity. The assembler output was polished one time using Racon^12^ with Nanopore reads, then one time with Medaka^13^ and Nanopore reads and two times with Hapo-G^14^ and Illumina PCR-free reads.(Table S2).

### Assembly validation

The DLE-1 map was generated using the Direct Label and Stain (DLS) technology and the BspQI map using the Nick Label Repair and Stain (NLRS) technology. Genome map assemblies were performed using Bionano Solve Pipeline version 3.3 and Bionano Access version 1.3.0. We used the parameter “Add Pre-Assembly” which produced a rough assembly. This first result was used as a reference for a second assembly, using the parameters “non-haplotype without extend and split”. We filtered out molecules smaller than 150 Kb and molecules with less than nine labelling sites (Tables S3 and S4). The nanopore contigs were then validated using the two Bionano maps and organized with the scaffolding procedure provided by Bionano Genomics (Figure S2). Negative gap sizes were checked and corrected using the BiscoT software^15^ to avoid artifactual genomic duplications (Table S6). As recommended by the BiscoT authors, we performed a last iteration of Hapo-G^14^ to polish merged regions (Table S5).

### Chromosomes reconstruction

DH-Pahang sequences were anchored on chromosomes using segregating markers obtained from the selfing of the ‘Pahang’ accession PT-BA-00267, described in Martin et al^10^ (Table S7 and Figure S3). Data are available on the Banana Genome Hub^44^ in the download section under ‘*AF-Pahang marker matrix file*’ and ‘*AF-Pahang marker sequence (FASTA)*’ for coded segregating markers and marker sequence respectively. Sequences anchoring was performed following methodology described in Martin et al.^10^. The complete process was performed using scaffhunter tools^45^ available at the South Green platform.

In addition, based on scaffold BLAST against *Musa acuminata* chloroplast sequence^46^, *Musa acuminata* putative 12 mitochondrial scaffolds^10^ and *Phoenix dactylifera* protein sequences^47^, 1 putative chloroplastic (corresponding to one initial contig) and 45 putative mitochondrial scaffolds (corresponding to 45 initial contigs) were identified in the assembly. The 45 putative mitochondrial scaffolds were grouped into a sequence named *putative_mito_scaff*. The chloroplastic scaffold was discarded from the assembly as the chloroplast genome of DH-Pahang was already fully assembled and published^46^.

The 37 remaining scaffolds (cumulative size of 5.2Mb) (corresponding each to one initial contig) showed a strong BLAST homology to larger scaffolds included in the chromosomes (36 with more than 95% of their length and 1 with more than 88% of its length). Investigation (dot-plot analysis using gepard v1.30^48^ and BLAST against nr/nt of ncbi) of these scaffolds revealed a repetitive nature, most of them corresponding to rDNA sequences. Because of their strong homology to scaffolds included in the chromosomes these scaffolds were discarded from the assembly.

### Gene prediction

Repeats in the genome assembly were masked using Tandem Repeat Finder^49^ for tandem repeats and RepeatMasker^50^ for simple repeats as well as known repeats included in RepBase^51^. In addition, known *Musa* transposable elements (from D’Hont et al^9^), were detected using RepeatMasker.

Gene prediction was done using proteomes from homologous species, *Musa acuminata* (UP000012960), *Oryza. sativa* (UP000059680), *Phoenix dactylifera* (UP000228380), *Musa schizocarpa* (www.genoscope.cns.fr/plants) and *Musa balbisiana* (banana-genome-hub.southgreen.fr).

The proteomes were aligned against the genome assembly in two steps. Firstly, BLAT^52^ (default parameters) was used to quickly localize corresponding putative genes of the proteins on the genome. The best match and matches with a score ≥ 90% of the best match score were retained. Secondly, the alignments were refined using Genewise^53^ (default parameters), which is more precise for intron/exon boundary detection. Alignments were kept if more than 80% of the length of the protein was aligned to the genome.

To allow the detection of UTRs in the gene prediction step, we aligned not only the protein of *M. acuminata*, but also the virtual mRNAs of the *M. acuminata* predicted genes^54^ on a masked version of the genome assembly. Then, transcript sequences of *M. acuminate* predicted genes were aligned by BLAT (default parameters) on the masked genome assembly. Only the alignments with an identity percent greater or equal to 90% were kept. For each transcript, the best match was selected based on the alignment score. Finally, alignments were recomputed in the previously identified genomic regions by Est2Genome^55^ in order to define precisely intron boundaries. Alignments were kept if more than 80% of the length of the transcript was aligned to the genome with a minimal identity percent of 95%.

To proceed to the gene prediction, we integrated the protein homologies and transcript mapping using a combiner called Gmove^56^. This tool can find CDSs based on genome located evidence without any calibration step. Briefly, putative exons and introns, extracted from the alignments, were used to build a simplified graph by removing redundancies. Then, Gmove extracted all paths from the graph and searched for open reading frames (ORFs) consistent with the protein evidence. A selection step was applied to all candidate genes, essentially based on gene structure. Also, all gene predictions included in a genomic region tagged as transposable elements were removed from the gene set (Table 2). Completeness of the predicted genes was assessed with BUSCO^57^ version 5 (embryophyta dataset odb10).

The search for NLR loci was performed using NLR-annotator tools^25^ that scan specifically the 6 reading frames of the nucleotide sequence for the presence of 19 NLR-associated motifs and reconstruct a potential NLR locus which might correspond to a complete or partial gene and might also be a pseudogene.

### Transposable element detection

Transposable elements were detected using RepeatMasker^50^ associated with the TE Musa library^9^ and CR sequences^17^. The same procedure was used to detect TEs in the Musa *acuminata* V2, V4, *Musa balbisiana* and *Musa schizocarpa* assemblies. The gff output file was converted into a bed file and the TE coverage was calculated using bedtools^58^ coverage (version v2.29.2-17-ga9dc5335) on a 100Kb window. Centromeric boundaries were defined using the density of daterra-Maximus, ITS-5S, ITS-18S, ITS-26S, Nanica, maca-Angela, caturra-Reina elements. TEs were grouped following Wicker et al. classification^59^.

### Genome comparisons

The synteny relationships between *Musa balbisiana, Musa schizocarpa* and *Musa acuminata* V4 were determined using Assemblytics^60^. First, genome sequences were aligned against each other using nucmer^61^ version 3.23 (-maxmatch -l 100 -c 500) as recommended by the Assemblytics authors. Assemblytics was launched on the nucmer delta file (unique_length_required = 10000). The figures 1 and 2B were generated using the Circos software^62^. In the same way, each chromosome of the *Musa acuminata* V4 was aligned against its relative chromosome of *Musa acuminata* V2 using nucmer version 3.23 (-r -1 -l 10000) and dotplots were generated using the mummerplot command.

### Detection of specific regions of the V4 assembly

New regions of the *Musa acuminata* V4 were determined using blast^63^ (ncbi-tools/6.1.20120620) alignment between the chromosomes of each assembly version. Regions of the V4 assembly larger than 100Kb without any alignment to the V2 assembly were considered new. In addition, the *Musa acuminata* V2 gene predictions were aligned (see Gene prediction section) on the V4 assembly and the positions of the genes were compared with the V4 gene catalogue using bedtools^58^ (version bedtools-2.29.2) (-v option). Genes from the V4 assembly without any correspondence in the V2 were considered new.

### Detection of tandemly duplicated genes

An all-against-all comparison of the *Musa acuminata* V4 proteins was performed using Diamond^64^ (version 0.9.24). Mapping output was filtered according to the following parameters : an e-value lower than 10e-20 and a coverage of the smallest protein greater than 80%. Genes were considered as tandemly duplicated if they were co-localized on the same chromosome and not distant of more than 10 genes to each other. Figure 4A was realized using the MCscan tool^65^ with the two following commands : jcvi.compara.synteny (--iter=1) and jcvi.graphics.synteny (--glyphcolor=orientation).

## Supporting information

Table S12 : Annotated NLR genes in the V4 assembly

Table S13 : Annotated NLR genes in the V2 assembly

## Additional files

All the supporting data are included in three additional files which contain a) Tables S1-S11 and Figures S1-S14, b) Table S12 (position of NLR genes in the V4 assembly) and c) Table S13 (position of NLR genes in the V2 assembly).

## Data availability

The genome assembly is freely available at http://www.genoscope.cns.fr/plants and http://banana-genome-hub.southgreen.fr. The ONT, Illumina and Bionano Genomics data are available in the European Nucleotide Archive under the following projects PRJEB35002.

## Acknowledgements

The authors thank the staff of Oxford Nanopore Technology Ltd for technical help, Jitka Weiserová, Eva Jahnová and Dr. Jan Vrána for their help with the material preparation, the CRB Plantes Tropicales Antilles CIRAD-INRA Guadeloupe France for providing the plant materials and the CIRAD – UMR AGAP HPC Data Center of the South Green Bioinformatics platform (http://www.southgreen.fr) for providing computational resources.

## Author’s information

KL extracted the sequenced DNA. EH and JD prepared HMW DNA for optical mapping. KL extracted the plugs. CC realized the bionano experiments. KL, CC and AL optimized and performed the sequencing. CB, BI, BN, NY, FCB, GM and JMA performed the bioinformatic analyses. AD and JMA conceived the project. CB, BI, BN, KL, CC, EH, GM, AD and JMA wrote the article. AD, PW and JMA supervised the study.

## Ethics declaration

JMA received travel and accommodation expenses to speak at Oxford Nanopore Technologies conferences. JMA and CB received accommodation expenses to speak during Bionano Genomics user meetings. The authors declare that they have no other competing interests.

## References

1. Michael, T. P. & VanBuren, R. Building near-complete plant genomes. Curr. Opin. Plant Biol. 54, 26–33 (2020).

2. Rousseau-Gueutin, M. et al. Long-read assembly of the Brassica napus reference genome Darmor-bzh. GigaScience 9, (2020).

3. Zhang, W. et al. Genome assembly of wild tea tree DASZ reveals pedigree and selection history of tea varieties. Nat. Commun. 11, 3719 (2020).

4. Schmidt, M. H.-W. et al. De Novo Assembly of a New Solanum pennellii Accession Using Nanopore Sequencing. Plant Cell 29, 2336–2348 (2017).

5. Miga, K. H. et al. Telomere-to-telomere assembly of a complete human X chromosome. Nature 585, 79–84 (2020).

6. Martin, G. et al. Genome ancestry mosaics reveal multiple and cryptic contributors to cultivated banana. Plant J. 102, 1008–1025 (2020).

7. Němečková, A. et al. Molecular and Cytogenetic Study of East African Highland Banana. Front. Plant Sci. 9, (2018).

8. Langhe, E. D., Vrydaghs, L., Maret, P. de Perrier, X. & Denham, T. Why Bananas Matter: An introduction to the history of banana domestication. Ethnobot. Res. Appl. 7, 165–177 (2009).

9. D’Hont, A. et al. The banana (Musa acuminata) genome and the evolution of monocotyledonous plants. Nature 488, 213–217 (2012).

10. Martin, G. et al. Improvement of the banana “Musa acuminata” reference sequence using NGS data and semi-automated bioinformatics methods. BMC Genomics 17, 243 (2016).

11. Chen, Y. et al. Efficient assembly of nanopore reads via highly accurate and intact error correction. Nat. Commun. 12, 60 (2021).

12. Vaser, R., Sović, I., Nagarajan, N. & Šikić, M. Fast and accurate de novo genome assembly from long uncorrected reads. Genome Res. 27, 737–746 (2017).

13. nanoporetech/medaka. (Oxford Nanopore Technologies, 2021).

14. Aury, J.-M. & Istace, B. Hapo-G, Haplotype-Aware Polishing Of Genome Assemblies. bioRxiv 2020.12.14.422624 (2020) doi:10.1101/2020.12.14.422624.

15. Istace, B., Belser, C. & Aury, J.-M. BiSCoT: improving large eukaryotic genome assemblies with optical maps. PeerJ 8, e10150 (2020).

16. Čížková, J. et al. Molecular Analysis and Genomic Organization of Major DNA Satellites in Banana (Musa spp.). PLOS ONE 8, e54808 (2013).

17. Tran, T. D. et al. Centromere and telomere sequence alterations reflect the rapid genome evolution within the carnivorous plant genus Genlisea. Plant J. Cell Mol. Biol. 84, 1087–1099 (2015).

18. Neumann, P. et al. Plant centromeric retrotransposons: a structural and cytogenetic perspective. Mob. DNA 2, 4 (2011).

19. Panchy, N., Lehti-Shiu, M. & Shiu, S.-H. Evolution of Gene Duplication in Plants. Plant Physiol. 171, 2294–2316 (2016).

20. Del Terra, L. et al. Functional characterization of three Coffea arabica L. monoterpene synthases: Insights into the enzymatic machinery of coffee aroma. Phytochemistry 89, 6–14 (2013).

21. Jiang, S.-Y., Jin, J., Sarojam, R. & Ramachandran, S. A Comprehensive Survey on the Terpene Synthase Gene Family Provides New Insight into Its Evolutionary Patterns. Genome Biol. Evol. 11, 2078–2098 (2019).

22. Falara, V. et al. The Tomato Terpene Synthase Gene Family. Plant Physiol. 157, 770–789 (2011).

23. Martin, D. M. et al. Functional Annotation, Genome Organization and Phylogeny of the Grapevine (Vitis vinifera) Terpene Synthase Gene Family Based on Genome Assembly, FLcDNA Cloning, and Enzyme Assays. BMC Plant Biol. 10, 226 (2010).

24. Wersch S. van & Li, X. Stronger When Together: Clustering of Plant NLR Disease resistance Genes. Trends Plant Sci. 24, 688–699 (2019).

25. Steuernagel, B. et al. The NLR-Annotator Tool Enables Annotation of the Intracellular Immune Receptor Repertoire. Plant Physiol. 183, 468–482 (2020).

26. Belser, C. et al. Chromosome-scale assemblies of plant genomes using nanopore long reads and optical maps. Nat. Plants 4, 879–887 (2018).

27. Wang, Z. et al. Musa balbisiana genome reveals subgenome evolution and functional divergence. Nat. Plants 5, 810–821 (2019).

28. Yang, X. et al. Amplification and adaptation of centromeric repeats in polyploid switchgrass species. New Phytol. 218, 1645–1657 (2018).

29. Miga, K. H. Centromere studies in the era of ‘telomere-to-telomere’ genomics. Exp. Cell Res. 394, 112127 (2020).

30. Comai, L., Maheshwari, S. & Marimuthu, M. P. A. Plant centromeres. Curr. Opin. Plant Biol. 36, 158–167 (2017).

31. Bellaire, L. de L. de Fouré, E., Abadie, C. & Carlier, J. Black Leaf Streak Disease is challenging the banana industry. Fruits 65, 327–342 (2010).

32. Kema, G. H. J. et al. Editorial: Fusarium Wilt of Banana, a Recurring Threat to Global Banana Production. Front. Plant Sci. 11, (2021).

33. Ahmad, F. et al. Genetic mapping of Fusarium wilt resistance in a wild banana Musa acuminata ssp. malaccensis accession. Theor. Appl. Genet. 133, 3409–3418 (2020).

34. Gawel, N. J. & Jarret, R. L. A modified CTAB DNA extraction procedure forMusa andIpomoea. Plant Mol. Biol. Report. 9, 262–266 (1991).

35. Safár, J. et al. Creation of a BAC resource to study the structure and evolution of the banana (Musa balbisiana) genome. Genome 47, 1182–1191 (2004).

36. Šimková, H., Číhalíková, J., Vrána, J., Lysák, M. A. & Doležel, J. Preparation of HMW DNA from Plant Nuclei and Chromosomes Isolated from Root Tips. Biol. Plant. 46, 369–373 (2003).

37. Engelen S, Aury JM. fastxtend. https://www.genoscope.cns.fr/externe/fastxtend/.

38. Li, R., Li, Y., Kristiansen, K. & Wang, J. SOAP: short oligonucleotide alignment program. Bioinformatics 24, 713–714 (2008).

39. Alberti, A. et al. Viral to metazoan marine plankton nucleotide sequences from the Tara Oceans expedition. Sci. Data 4, 170093 (2017).

40. rrwick/Filtlong: quality filtering tool for long reads. https://github.com/rrwick/Filtlong.

41. Liu, H., Wu, S., Li, A. & Ruan, J. SMARTdenovo: a de novo assembler using long noisy reads. Gigabyte 2021, 1–9 (2021).

42. Ruan, J. & Li, H. Fast and accurate long-read assembly with wtdbg2. Nat. Methods 17, 155–158 (2020).

43. Kolmogorov, M., Yuan, J., Lin, Y. & Pevzner, P. A. Assembly of long, error-prone reads using repeat graphs. Nat. Biotechnol. 37, 540–546 (2019).

44. Droc, G. et al. The Banana Genome Hub. Database 2013, (2013).

45. SouthGreenPlatform/scaffhunter. (South Green Bioinformatics platform, 2019).

46. Martin, G., Baurens, F.-C., Cardi, C., Aury, J.-M. & D’Hont, A. The Complete Chloroplast Genome of Banana (Musa acuminata, Zingiberales): Insight into Plastid Monocotyledon Evolution. PLOS ONE 8, e67350 (2013).

47. Fang, Y. et al. A Complete Sequence and Transcriptomic Analyses of Date Palm (Phoenix dactylifera L.) Mitochondrial Genome. PLOS ONE 7, e37164 (2012).

48. Krumsiek, J., Arnold, R. & Rattei, T. Gepard: a rapid and sensitive tool for creating dotplots on genome scale. Bioinformatics 23, 1026–1028 (2007).

49. Benson, G. Tandem repeats finder: a program to analyze DNA sequences. Nucleic Acids Res. 27, 573–580 (1999).

50. Smit, AFA, Hubley, R & Green, P. RepeatMasker. http://repeatmasker.org/.

51. Bao, W., Kojima, K. K. & Kohany, O. Repbase Update, a database of repetitive elements in eukaryotic genomes. Mob. DNA 6, 11 (2015).

52. Kent, W. J. BLAT—The BLAST-Like Alignment Tool. Genome Res. 12, 656–664 (2002).

53. Birney, E., Clamp, M. & Durbin, R. GeneWise and Genomewise. Genome Res. 14, 988 (2004).

54. Improvement of the banana “ Musa acuminata “ reference sequence using NGS data and semi-automated bioinformatics methods | BMC Genomics | Full Text. https://bmcgenomics.biomedcentral.com/articles/10.1186/s12864-016-2579-4.

55. Mott, R. EST_GENOME: a program to align spliced DNA sequences to unspliced genomic DNA. Comput. Appl. Biosci. CABIOS 13, 477–478 (1997).

56. Dubarry, M. et al. Gmove a tool for eukaryotic gene predictions using various evidences. F1000Research 5, (2016).

57. Waterhouse, R. M. et al. BUSCO Applications from Quality Assessments to Gene Prediction and Phylogenomics. Mol. Biol. Evol. 35, 543–548 (2018).

58. Quinlan, A. R. & Hall, I. M. BEDTools: a flexible suite of utilities for comparing genomic features. Bioinformatics 26, 841–842 (2010).

59. Wicker, T. et al. A unified classification system for eukaryotic transposable elements. Nat. Rev. Genet. 8, 973–982 (2007).

60. Nattestad, M. & Schatz, M. C. Assemblytics: a web analytics tool for the detection of variants from an assembly. Bioinformatics 32, 3021–3023 (2016).

61. Kurtz, S. et al. Versatile and open software for comparing large genomes. Genome Biol. 5, R12 (2004).

62. Krzywinski, M. I. et al. Circos: An information aesthetic for comparative genomics. Genome Res. (2009) doi:10.1101/gr.092759.109.

63. Altschul, S. F., Gish, W., Miller, W., Myers, E. W. & Lipman, D. J. Basic local alignment search tool. J. Mol. Biol. 215, 403–410 (1990).

64. Buchfink, B., Xie, C. & Huson, D. H. Fast and sensitive protein alignment using DIAMOND. Nat. Methods 12, 59–60 (2015).

65. Tang, H. et al. Synteny and Collinearity in Plant Genomes. Science 320, 486–488 (2008).

