## Supplementary Data for "Telomere-to-telomere gapless chromosomes of banana using nanopore sequencing"

**Table S1: Statistics of the ONT datasets.**

| | DH-Pahang | | |
|---|---|---|---|
| | Raw reads PromethION | Longest reads | Filtlong reads |
| Cumulative size | 92,749,841,432 | 13,500,017,777 | 13,500,035,543 |
| # of reads | 5,185,398 | 175,937 | 273,413 |
| Coverage | 206x | 30x | 30x |
| N50 (bp) | 31,640 | 74,681 | 52,825 |

**Table S2: DH-Pahang ONT assembly statistics**.

| | SMARTDeNovo | | | Redbean | | |
|---|---|---|---|---|---|---|
| Subset of reads used | All reads | Longest | Filtlong | All reads | Longest | Filtlong |
| Cumulative size | 482,349,453 | 465,668,591 | 431,317,546 | 2,021,646,269 | 802,627,210 | 606,906,240 |
| # contigs | 121 | 183 | 783 | 43,296 | 13,785 | 10,307 |
| N50 (L50) | 19,507,988 (9) | 5,667,738 (22) | 977,146 (111) | 75,016 (6,675) | 93,433 (1,502) | 92,564 (989) |
| N90 (L90) | 2,709,408 (28) | 1,176,439 (92) | 230,199 (490) | 21,284 (26,326) | 25,463 (7,959) | 25,353 (6,103) |

| | Flye | | | NECAT |
|---|---|---|---|---|
| Subset of reads used | All reads | Longest | Filtlong | All reads |
| Cumulative size | 470,287,666 | 471,731,899 | 439,685,078 | 485,698,222 |
| # contigs | 449 | 318 | 953 | 175 |
| N50 (L50) | 15,404,016 (12) | 12,916,122 (13) | 1,521,384 (73) | 32,031,704 (7) |
| N90 (L90) | 2,621,850 (40) | 3,315,873 (37) | 298,601 (333) | 5,650,309 (17) |

### Table S3: DH-Pahang bionano dataset

| | DLE-1 Molecule statistics | BspQI Molecule statistics |
|---|---|---|
| Total number of molecules | 2,151,406 | 2,897,832 |
| Total length (Mbp) | 209,178.556 | 398,318,111 |
| Average length (kbp) | 97.229 | 137.109 |
| Molecule N50 (kbp) | 129.750 | 220.875 |
| Label density (/100kb) | 13.709 | 10.502 |
| Number of Flow cell | 2 | 1 |

### Table S4: DH-Pahang bionano genome map

| | DLE-1 genome map | BspQI genome map |
|---|---|---|
| Genome map number | 24 | 71 |
| Total Genome Map Length (Mbp) | 469.764 | 474.019 |
| Genome Map N50 (Mbp) | 35.022 | 16.002 |

### Table S5: DH-Pahang hybrid scaffolding and polishing

| | nanopore contigs polished | Hybrid scaffolds | contigs not scaffolded | final hybrid scaffolds (hybrid scaffolds + contigs not scaffolded) | scaffolds after negative gap resolution and polishing |
|---|---|---|---|---|---|
| number | 124 | 16 | 80 | 96 | 97 |
| N50 (L50) | 32,091,274 (7) | 39,508,388 (6) | 249,463 (18) | 39,508,388 (6) | 39,373,400 (6) |
| N90 (L90) | 5,668,018 (17) | 21,536,064 (12) | 87,718 (59) | 21,536,064 (12) | 21,536,112 (12) |
| maxSize | 47,719,325 | 47,719,325 | 673,878 | 47,719,325 | 47,719,527 |
| Assembly size | 485,318,484 | 471,709,278 | 14,435,609 | 486,144,887 | 484,747,212 |
| % of N | 0% | 0.18% | 0% | 0.17% | 0.14% |

**Table S6: Contigs details before and after negative gap resolution.** Gaps of 100bp are gaps of unknown length generated by the anchoring of contigs using the genetic map. Gaps of 13bp are gaps of unknown size generated by the BioNano pipeline.

| DH-Pahang chromosome | # corresponding NECAT contigs | # hybrid scaffolds | # contigs after negative gap resolution | Gaps length (bp) |
|---|---|---|---|---|
| chr01 | 7 | 1 | 5 | 163,709 - 13 - 37,440 - 13 |
| chr02 | 1 | 1 | 1 | / |
| chr03 | 4 | 2 | 2 | 100 |
| chr04 | 1 | 1 | 1 | / |
| chr05 | 4 | 1 | 4 | 53,161 - 30,868 - 32,283 |
| chr06 | 1 | 1 | 1 | / |
| chr07 | 5 | 1 | 4 | 13 - 13 - 13 |
| chr08 | 4 | 2 | 4 | 100 - 46,534 - 324,396 |
| chr09 | 1 | 1 | 1 | / |
| chr10 | 8 | 2 | 2 | 100 |
| chr11 | 1 | 1 | 1 | / |

**Table S7**: **Marker number per scaffold and linkage group**

| scaffold | LG01-04 | LG02 | LG03 | LG05 | LG06 | LG07 | LG08 | LG09 | LG10 | LG11 |
|---|---|---|---|---|---|---|---|---|---|---|
| Pahang_Scaffold_1 | 0 | 0 | 0 | 1 | 0 | 1 | 0 | 2089 | 0 | 0 |
| Pahang_Scaffold_2 | 0 | 0 | 0 | 1874 | 0 | 1 | 0 | 0 | 1 | 0 |
| Pahang_Scaffold_3 | 2191 | 1 | 0 | 0 | 0 | 0 | 0 | 1 | 0 | 0 |
| Pahang_Scaffold_4 | 0 | 0 | 1 | 0 | 2190 | 0 | 0 | 1 | 0 | 0 |
| Pahang_Scaffold_5 | 1082 | 0 | 2 | 3 | 0 | 0 | 2 | 1 | 0 | 0 |
| Pahang_Scaffold_6 | 0 | 0 | 0 | 1 | 0 | 1687 | 0 | 0 | 1 | 1 |
| Pahang_Scaffold_7 | 0 | 0 | 0 | 0 | 1 | 0 | 0 | 0 | 1784 | 0 |
| Pahang_Scaffold_8 | 0 | 1483 | 0 | 0 | 0 | 0 | 0 | 0 | 0 | 0 |
| Pahang_Scaffold_9 | 0 | 0 | 0 | 1 | 0 | 0 | 1 | 1 | 0 | 1500 |
| Pahang_Scaffold_10 | 0 | 0 | 0 | 1 | 0 | 0 | 1283 | 0 | 0 | 0 |
| Pahang_Scaffold_11 | 0 | 0 | 1027 | 1 | 0 | 0 | 0 | 0 | 0 | 0 |
| Pahang_Scaffold_12 | 0 | 0 | 0 | 0 | 0 | 0 | 1276 | 0 | 0 | 0 |
| Pahang_Scaffold_13 | 0 | 0 | 850 | 0 | 0 | 0 | 1 | 0 | 1 | 0 |
| Pahang_Scaffold_14 | 0 | 0 | 0 | 1 | 0 | 0 | 0 | 0 | 173 | 0 |
| Pahang_Scaffold_59 | 1 | 0 | 0 | 0 | 0 | 0 | 0 | 0 | 0 | 0 |

**Table S8: TE classes proportions in *Musa acuminata* (V2 and V4), *Musa schizocarpa* and *Musa balbisiana*.**

| | | *Musa acuminata* V4 (%) | *Musa acuminata* V2 (%) | *Musa schizocarpa* (%) | *Musa balbisiana* (%) |
|---|---|---|---|---|---|
| Class I (retrotransposons) | | | | | |
| LTR | Copia | 11,079 | 8,965 | 13,766 | 12,432 |
| | Gypsy | 5,986 | 4,668 | 5,848 | 5,137 |
| | no cat | 17,788 | 12,743 | 19,767 | 16,547 |
| DIRS | RYX | 6,323 | 3,252 | 6,094 | 4,157 |
| PLE | Penelope | 0,003 | 0,003 | 0,003 | 0,004 |
| LINE | RIL/RIX | 3,492 | 2,741 | 3,478 | 2,941 |
| SINE | RSX | 0,005 | 0,006 | 0,009 | 0,004 |
| Large Retro-transposon Derivatives | RXX | 2,678 | 1,347 | 2,590 | 1,597 |
| Class II (DNA transposons)-Subclass 1 | | | | | |
| TIR | DTX | 0,163 | 0,172 | 0,180 | 0,185 |
| hAT | DTA | 0,506 | 0,538 | 0,525 | 0,637 |
| Class II (DNA transposons)-Subclass 2 | | | | | |
| Helitron | DHH/DHX | 2,292 | 2,175 | 1,737 | 2,152 |
| Maverick | DMX | 0,005 | 0,006 | 0,007 | 0,006 |
| MITE (miniature inverted repeat transposable elements) | DXX | 0,029 | 0,028 | 0,023 | 0,027 |
| No categories | | 0,722 | 0,549 | 1,097 | 0,762 |
| simple repeat | | 1,549 | 1,190 | 1,221 | 2,767 |
| | TOTAL | 52,623 | 38,383 | 56,345 | 49,353 |

**Table S9: Proportion of new annotated genes in each *Musa acuminata* chromosome of the V4 assembly**.

| | Nb of annotated genes | Nb of tandemly duplicated genes | Nb of new annotated genes | Nb of new tandemly duplicated genes |
|---|---|---|---|---|
| chr01 | 2,757 | 397 (14.40%) | 369 (13.38%) | 173 (46,88%) |
| chr02 | 2,680 | 300 (11.19%) | 232 (8.66%) | 88 (37,93%) |
| chr03 | 3,533 | 314 (8.89%) | 235 (6.65%) | 85 (36,17%) |
| chr04 | 4,254 | 343 (8.06%) | 258 (6.06%) | 66 (25,58%) |
| chr05 | 3,345 | 263 (7.86%) | 251 (7.50%) | 73 (29,08%) |
| chr06 | 4,076 | 349 (8.56%) | 283 (6.94%) | 96 (33,92%) |
| chr07 | 3,080 | 324 (10.52%) | 190 (6.17%) | 83 (43,68%) |
| chr08 | 3,604 | 299 (8.30%) | 221 (6.13%) | 67 (30,32%) |
| chr09 | 3,342 | 342 (10.23%) | 243 (7.27%) | 84 (34,57%) |
| chr10 | 3,552 | 527 (14.84%) | 510 (14.36%) | 250 (49,02%) |
| chr11 | 2,612 | 175 (6.70%) | 171 (6.55%) | 42 (24,56%) |
| Total | 36,835 | 3,633 (9.86%) | 2,963 (8.04%) | 1,137 (38.37%) |

**Table S10: Comparison of 5S ribosomal gene clusters in V2 and V4 Musa assemblies**

| | Base pair covered in V2 assembly (predicted genes) | Base pair covered in V4 assembly (predicted genes) |
|---|---|---|
| chr01 | 0 | 59,949 (1,882) |
| chr02 | 0 | 0 |
| chr03 | 194 (3) | 179,282 (4,645) |
| chr04 | 0 | 413 (10) |
| chr05 | 529 (21) | 640 (19) |
| chr06 | 68 (1) | 68 (1) |
| chr07 | 419 (7) | 216 (3) |
| chr08 | 4,104 (38) | 127,057 (1,135) |
| chr09 | 547 (14) | 80 (1) |
| chr10 | 116 (6) | 0 |
| chr11 | 0 | 0 |
| Un | 1,605 (40) | - |
| Total | 7,582 (130) | 367,705 (7,696) |

**Table S11: Comparison of NLR clusters between the V2 and V4 assemblies (based on NLR-Annotator predictions)**

| Assembly version | Chromosome | Cluster start coordinate | Cluster end coordinate | Size (bp) | Nb. Undetermined nucleotides (N) | Nb. NLR loci detected |
|---|---|---|---|---|---|---|
| V2 | chr03 | 27,854,779 | 27,992,717 | 137,938 | 5,255 | 14 |
| V2 | chr03 | 31,895,636 | 32,022,409 | 126,773 | 10,472 | 17 |
| V2 | chr07 | 29,765,171 | 29,842,567 | 77,396 | 4,269 | 6 |
| V2 | chr10 | 22,396,059 | 22,566,146 | 170,087 | 13,045 | 9 |
| V4 | chr03 | 36,566,894 | 36,731,875 | 164,981 | 0 | 16 |
| V4 | chr03 | 40,651,606 | 40,784,397 | 132,791 | 0 | 18 |
| V4 | chr07 | 33,870,053 | 34,018,178 | 148,125 | 0 | 10 |
| V4 | chr10 | 24,960,703 | 25,188,409 | 227,706 | 0 | 13 |

**Figure S1: gDNA extraction of DH-Pahang.** DNA quality was checked on a 2200 TapeStation automated electrophoresis system (Agilent, CA, USA). Ninety seven % of the DNA fragments have a length >50Kb. (a) before removal of small DNA fragments with Short Read Eliminator XL (Circulomics, MD, USA) (b) after removal of small DNA fragments

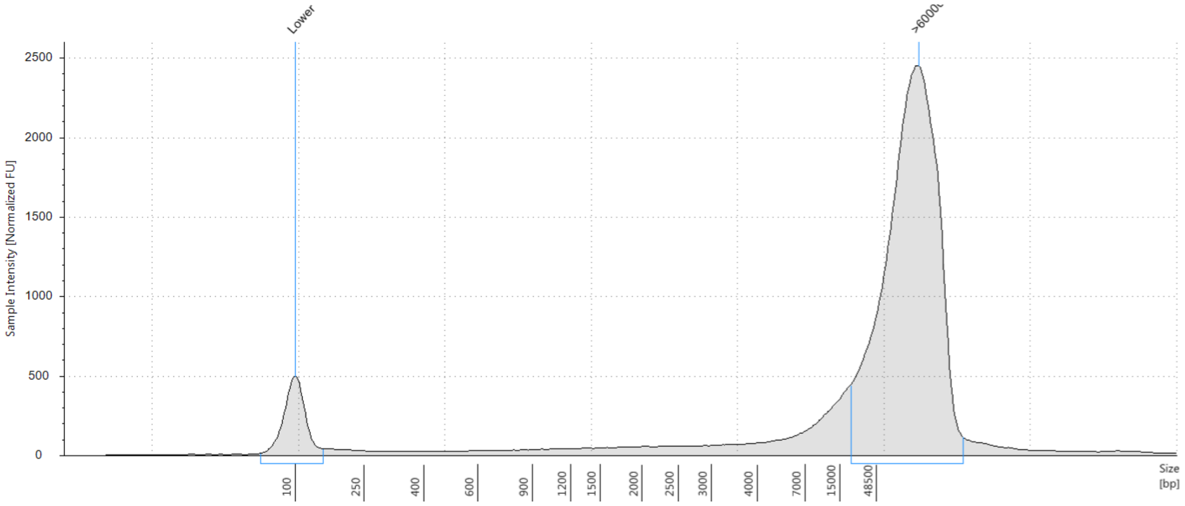


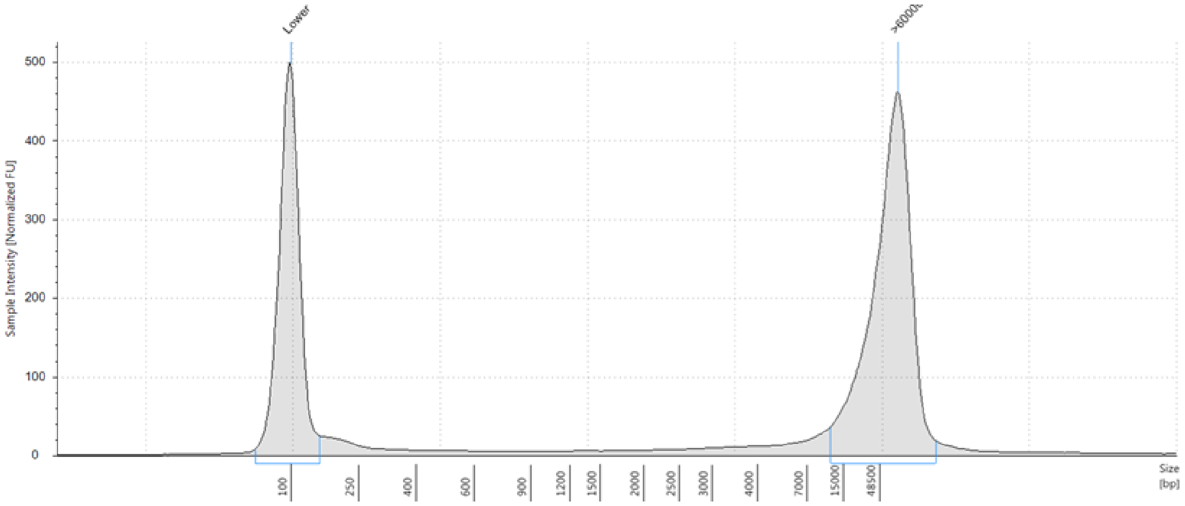

**Figure S2: Overview of the nanopore contigs in the V4 assembly.** Alignment of the two optical maps (a, red) DLE-1 and (b, red) BspQI with the nanopore contigs (c, blue). The vertical grey lines between the contigs and the optical maps represent the matches between enzymatic cuts from the map and the DNA sequences. The number of contigs of each chromosome is indicated.

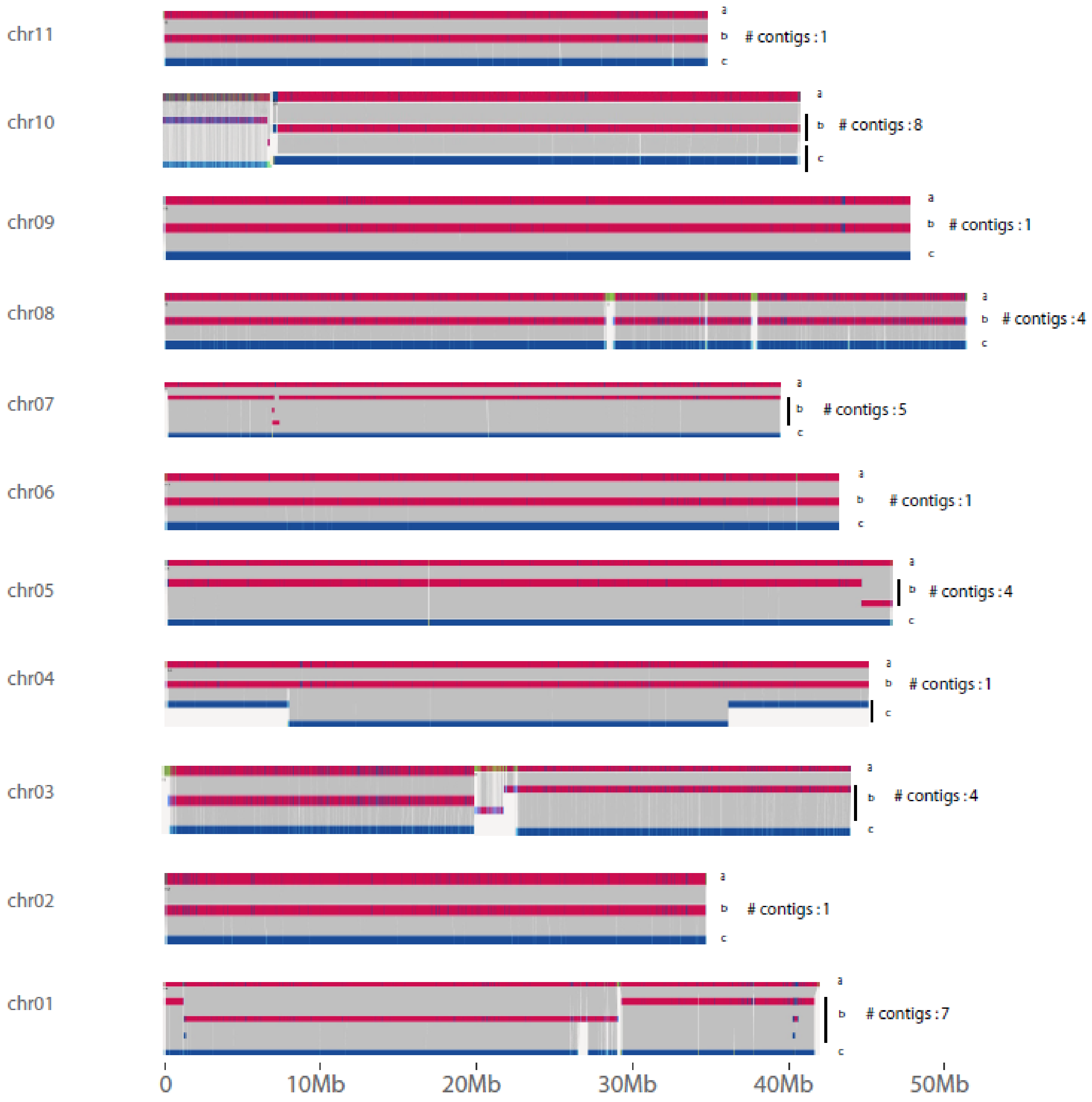

**Figure S3:** Dot plot showing marker linkage along ordered scaffolds of linkage group 01-04. This figure showed marker linkage of linkage group 01-04 which contained markers from chromosome 01 and 04. Because of chromosomal co-segregation due to reciprocal translocation between chromosome 01 and 04, markers from these chromosomes are linked. This resulted by the linkage of markers from a region of scaffold_3 to a region of scaffold_5. Interpretation of the figure suggested that in fact scaffold_3 corresponded to one chromosome and scaffold_5 corresponded to another chromosome.

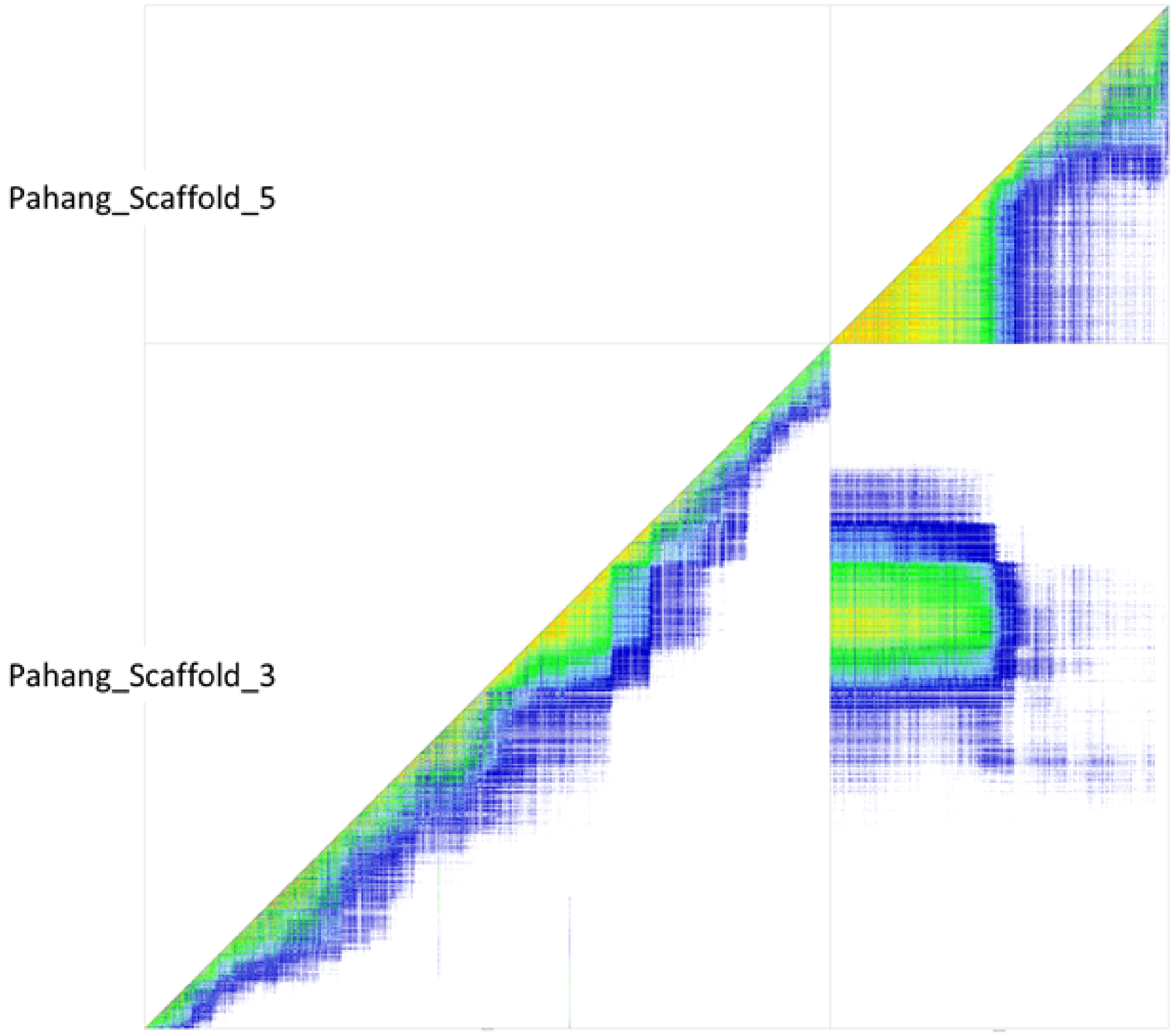

**Figure S4**: KAT plot of *Musa acuminata* V4 assembly.

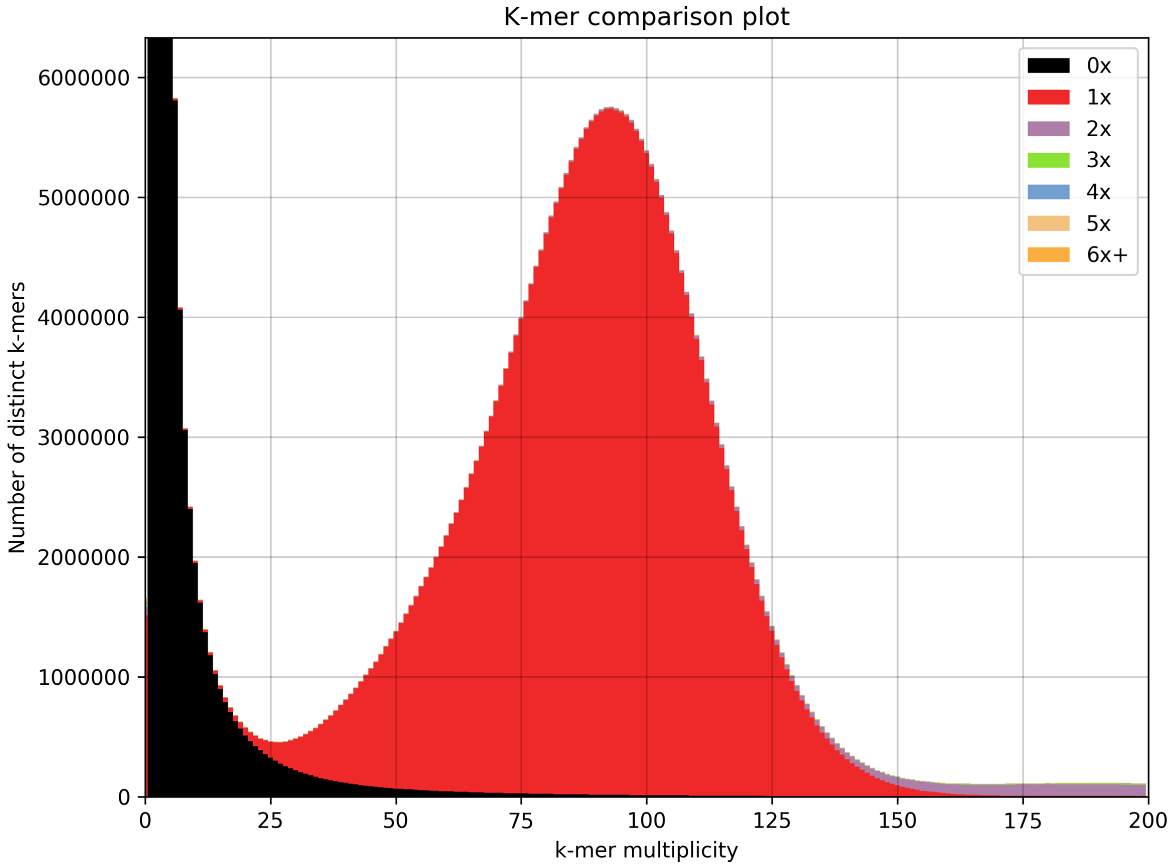

**Figure S5: Dot plot of each V4 chromosome against its relative V2 chromosome.** The x axis represents the chromosome in V2. The y axis represents the chromosome in V4. The number of the chromosome is printed in brackets.

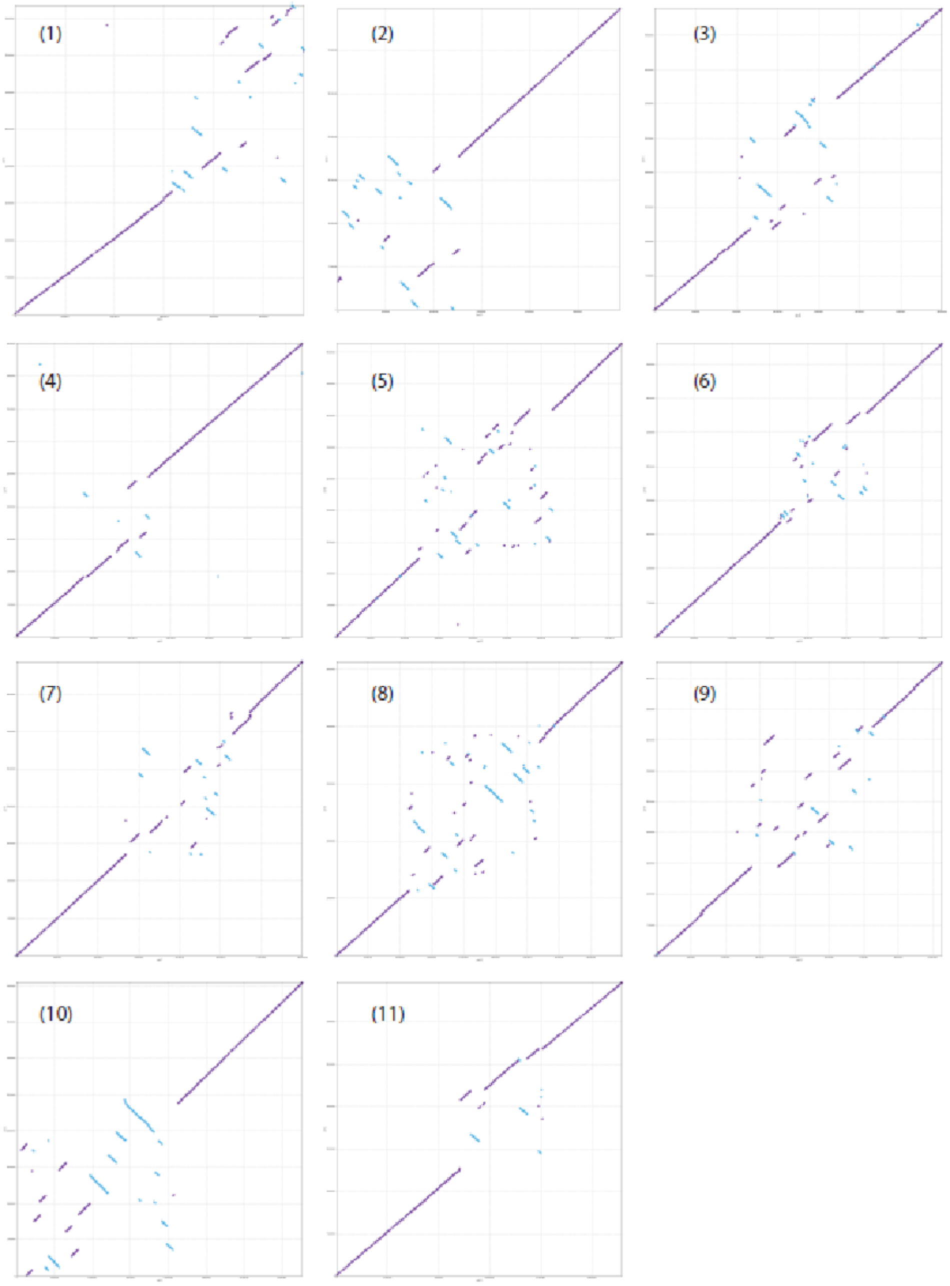

**Figure S6: Contig flagged as conflictual with the optical maps.** The contigs NGS41 (380,641bp) contains in tandem repetitive elements. The contig was split at the position 299,945b in two contigs of 299,945 et 80,696bp.

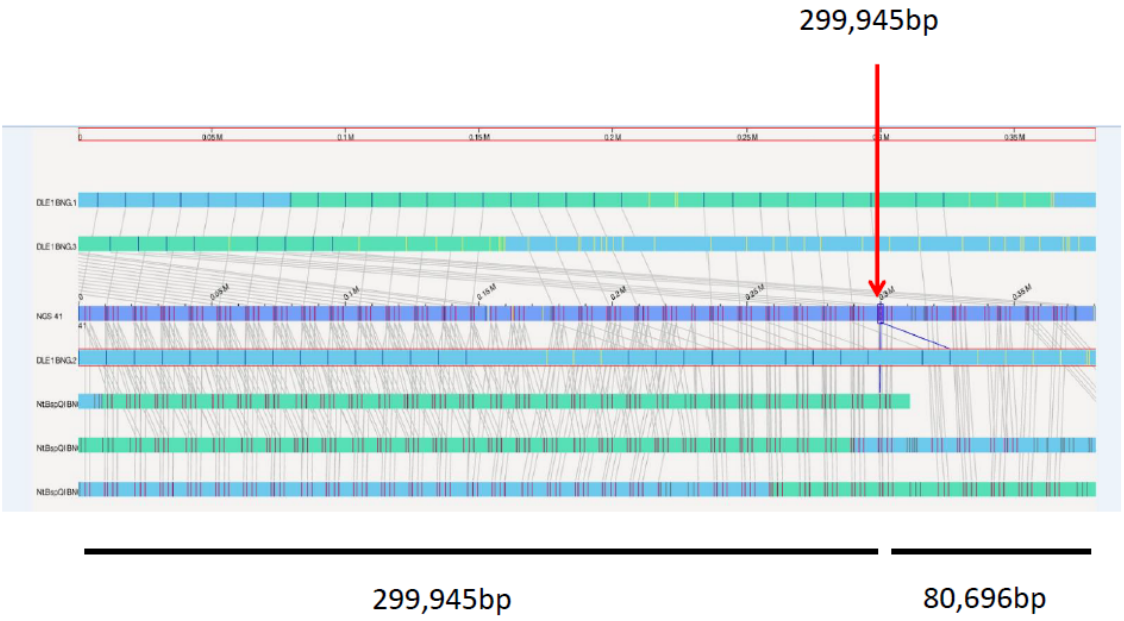

**Figure S7: Screenshot of the genome browser focusing on a *Musa acuminata* V4 chromosome 1 region.** These new annotated genes in the center of track 1 (Gene Predictions) are included in TDGs cluster and were absent from the *Musa acuminata* V2 annotation.

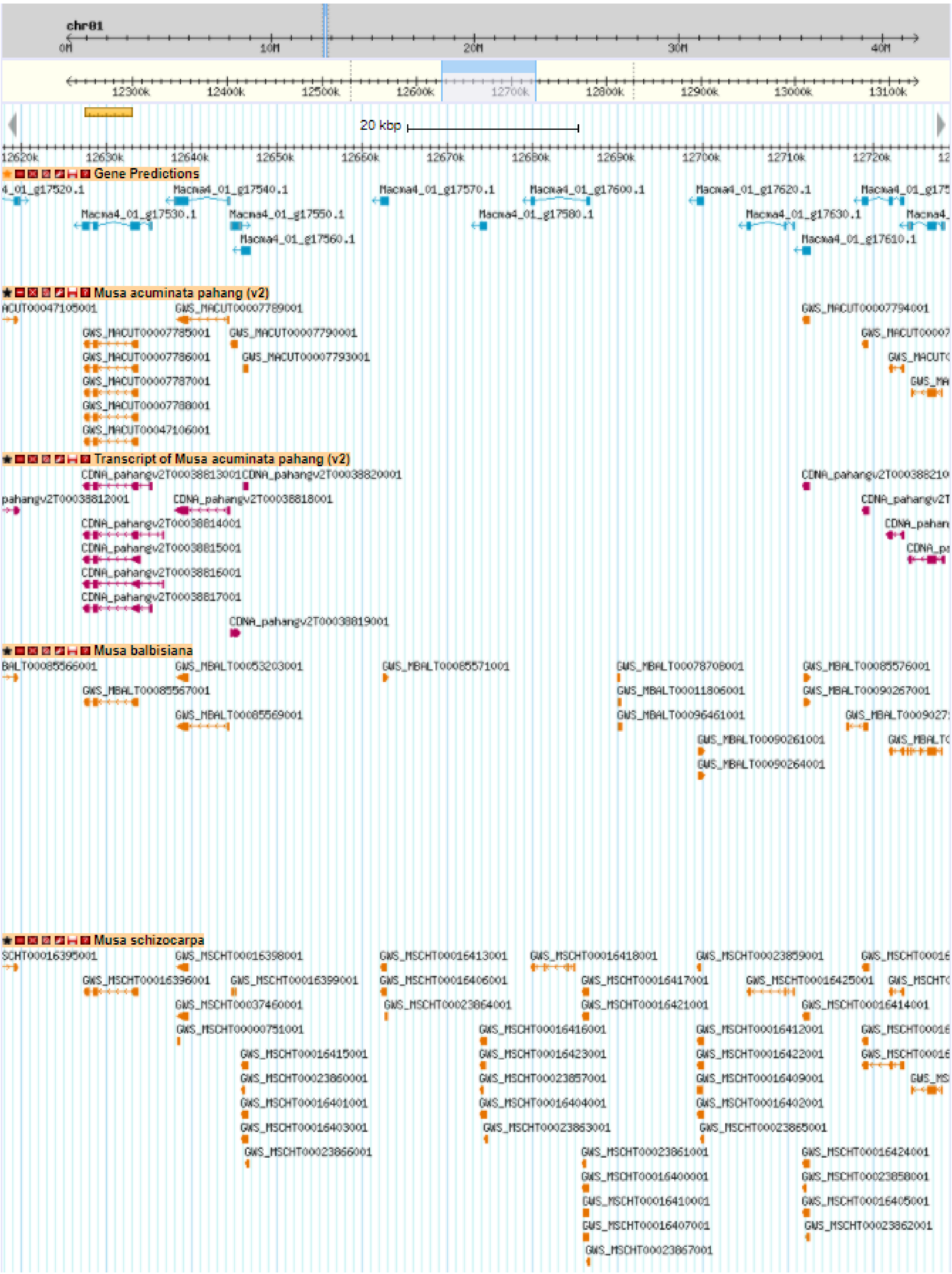

### Figure S8: Distribution of the repeats along the V4 chromosomes.

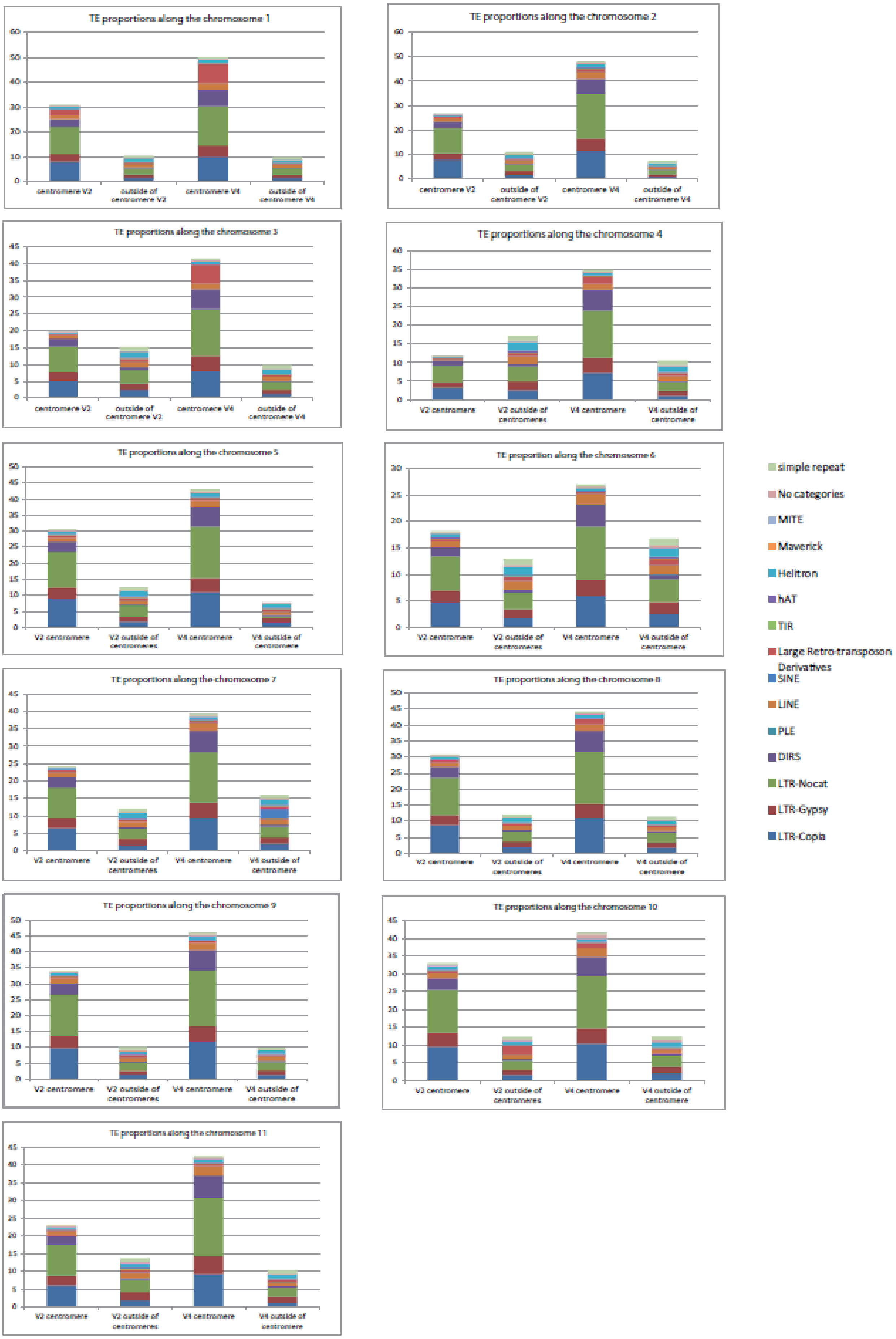

**Figure S9: Characterization of specific regions of the *Musa acuminata* V4 assembly.**

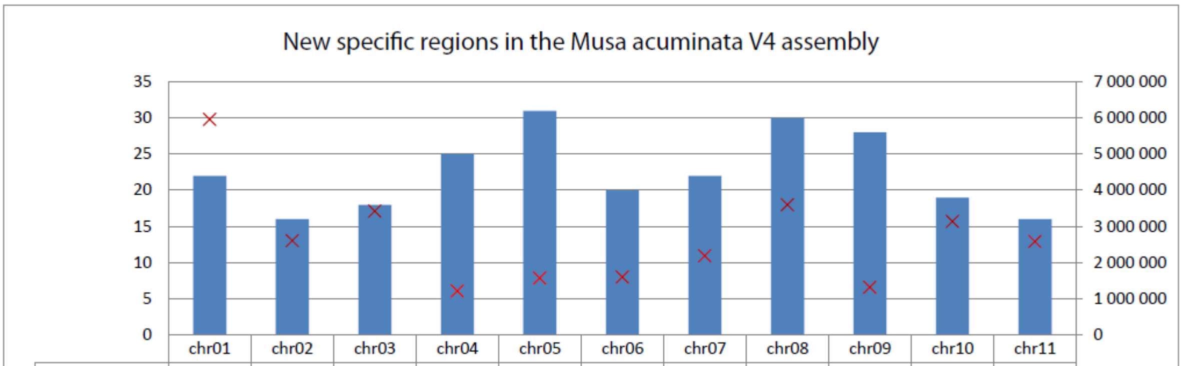


| | chr01 | chr02 | chr03 | chr04 | chr05 | chr06 | chr07 | chr08 | chr09 | chr10 | chr11 |
|---|---|---|---|---|---|---|---|---|---|---|---|
| # regions | 22 | 16 | 18 | 25 | 31 | 20 | 22 | 30 | 28 | 19 | 16 |
| Min length | 104 677 | 131 507 | 103 004 | 109 858 | 100 722 | 103 671 | 108 407 | 104 090 | 104 662 | 102 165 | 112 518 |
| Median length | 267 426 | 401 511 | 167 786 | 173 681 | 258 807 | 396 872 | 165 369 | 256 237 | 261 842 | 194 117 | 251 741 |
| Max length | 5 957 420 | 2 603 751 | 3 417 257 | 1 206 357 | 1 565 261 | 1 598 404 | 2 182 028 | 3 598 373 | 1 311 491 | 3 134 624 | 2 578 132 |

**Figure S10: Composition of specific regions of the *Musa acuminata* V4 assembly.**

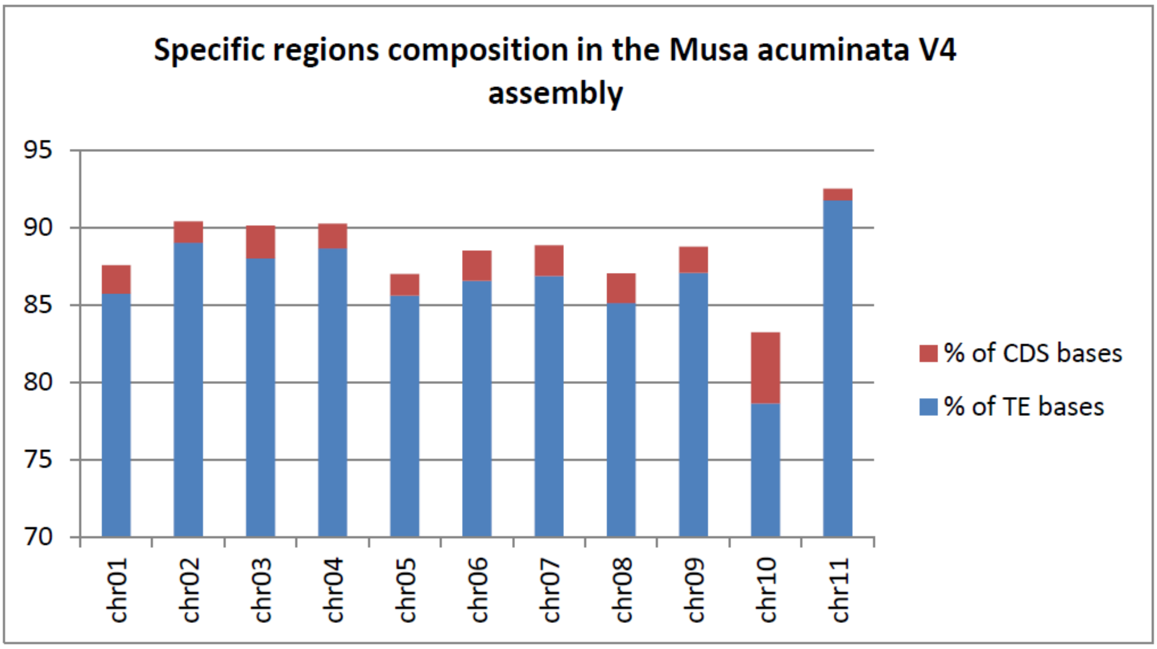

**Figure S11: Dot plot of *Musa balbisiana* assembly against *Musa acuminata* V4 assembly.**

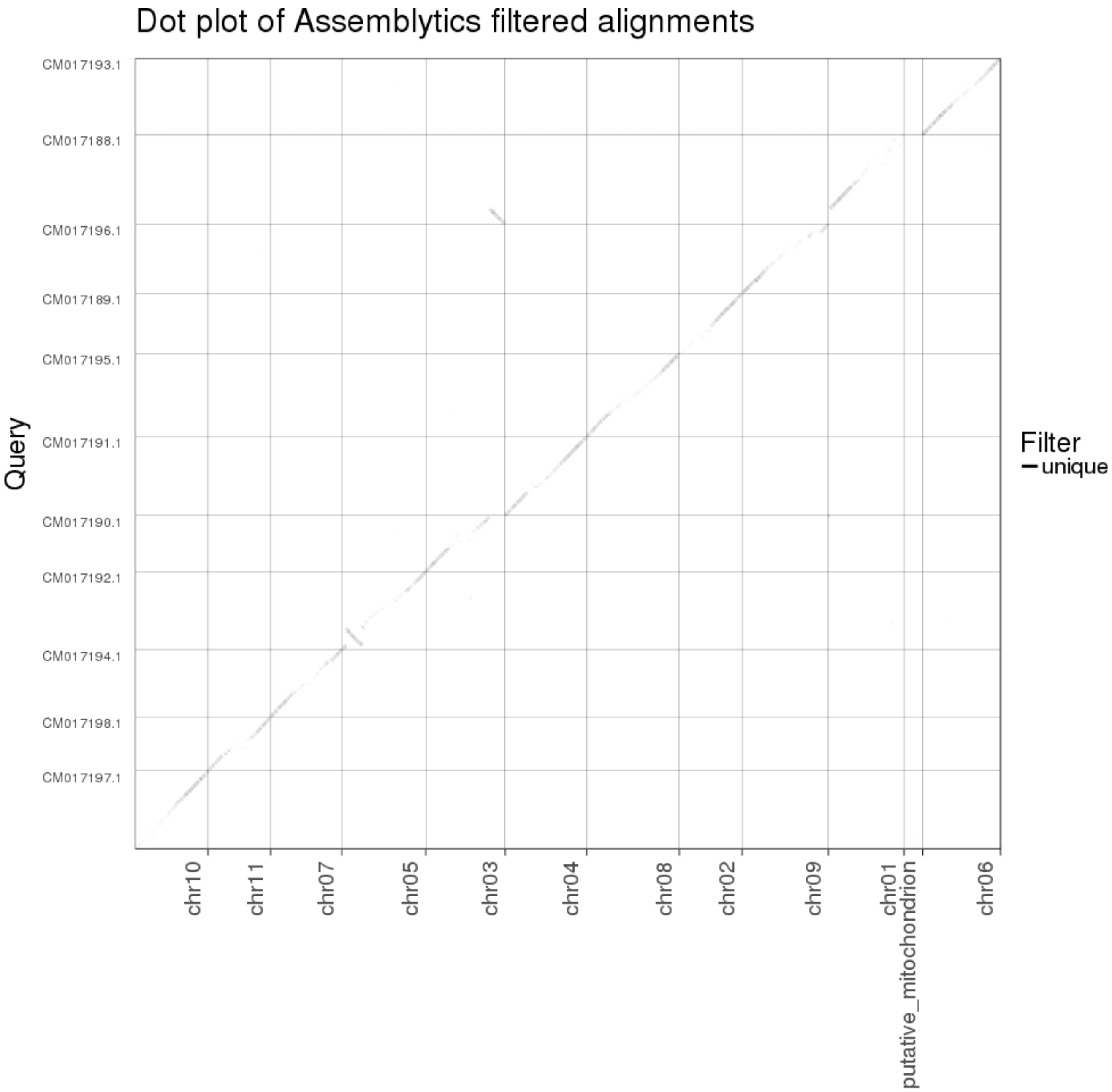

**Figure S12: Dot plot of *Musa schizocarpa* assembly against *Musa acuminata* V4 assembly**.

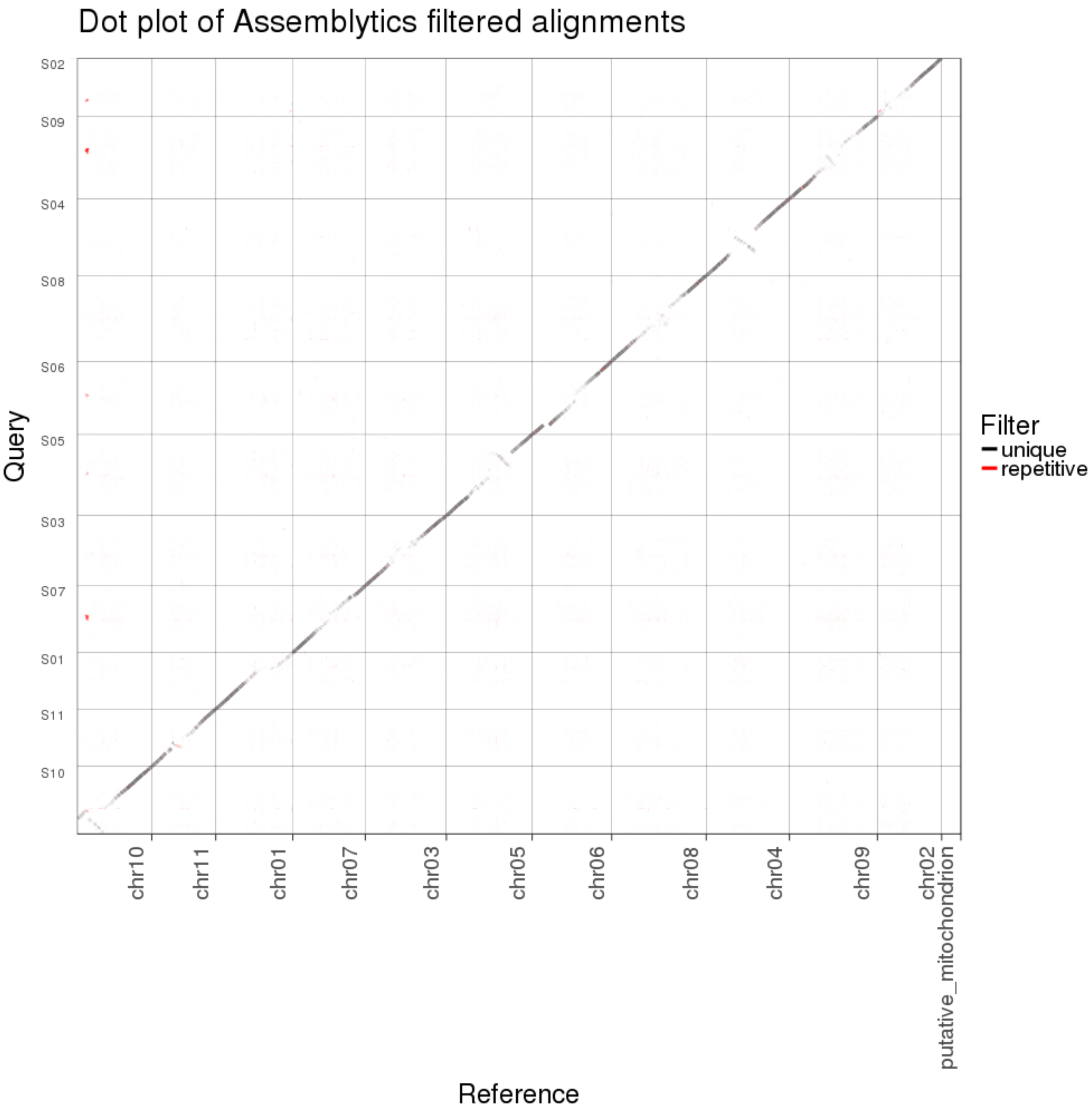

**Figure S13: Comparison of the structure of NLR loci clusters between DH-Pahang V2 and V4 assemblies.** The four panels represent dot plots of NLR loci clusters on chromosomes 3, 7 and 10 as indicated on top of each panel. The predicted NLR loci for each version are represented on the right side and at the bottom of the dot plots by blue boxes. Red boxes represent regions bearing undetermined nucleotides. Region coordinates are also indicated.

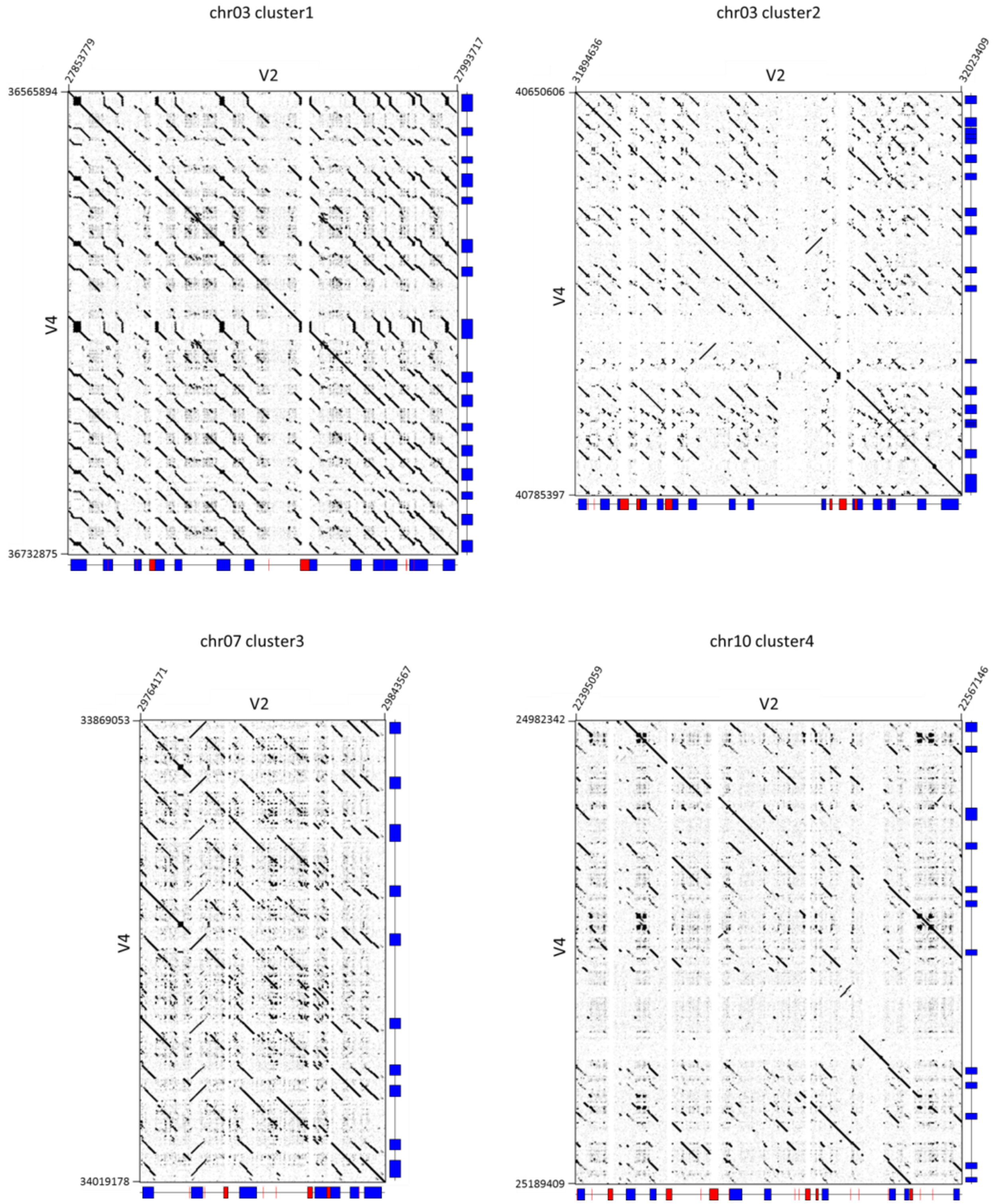

**Figure S14: Remaining gaps after negative gap resolution.** Example of a remaining sized gap in chromosome 1, located between the position 29,171,567 and 29,335,274. The gap was sized thanks to the BspQI map.

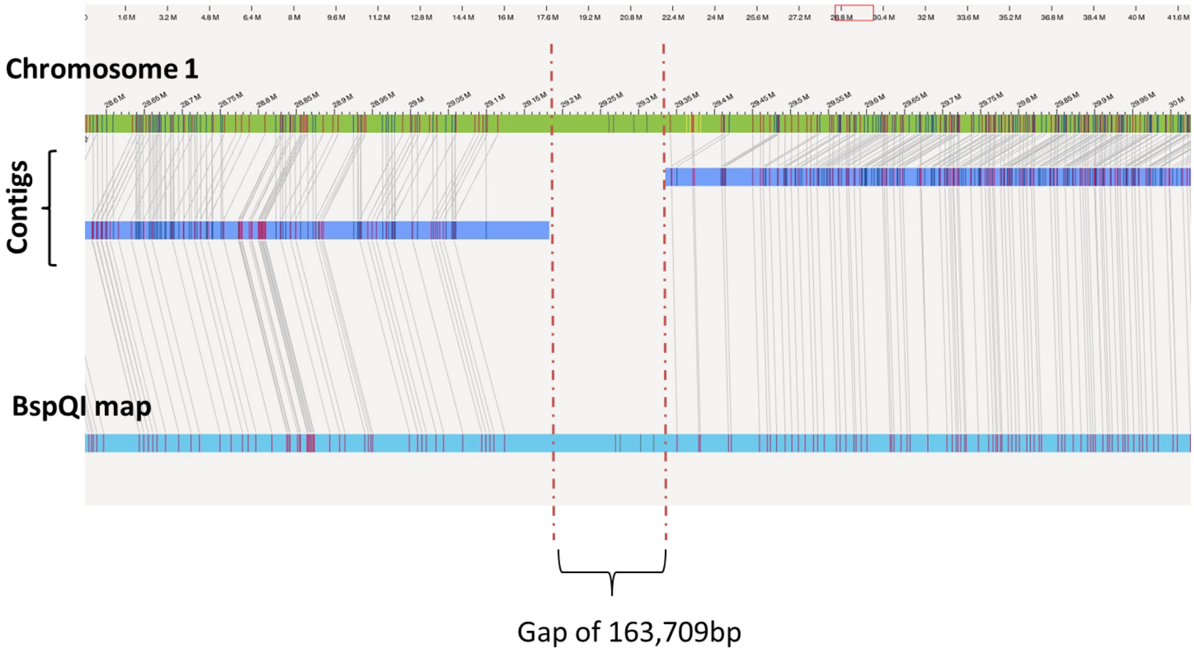
